# Selective ablation of adult GFAP-expressing tanycytes leads to hypogonadotropic hypogonadism in males

**DOI:** 10.1101/2021.07.31.454492

**Authors:** Lucile Butruille, Martine Batailler, Marie-Line Cateau, Ariane Sharif, Valérie Leysen, Vincent Prévot, Pascal Vaudin, Delphine Pillon, Martine Migaud

## Abstract

In adult mammals, neural stem cells emerge in three neurogenic regions, the subventricular zone of the lateral ventricle (SVZ), the subgranular zone of the dentate gyrus of the hippocampus (SGZ) and the hypothalamus. In the SVZ and the SGZ, neural stem/progenitor cells (NSPCs) express the glial fibrillary acidic protein (GFAP) and selective ablation of these NSPCs drastically decreases cell proliferation *in vitro* and *in vivo*. In the hypothalamus, GFAP is expressed by α-tanycytes, which are specialized radial glia-like cells in the wall of the third ventricle. To explore the role of these hypothalamic GFAP-positive tanycytes, we used transgenic mice expressing herpes simplex virus thymidine kinase (HSV-Tk) under the control of the mouse *Gfap* promoter and 4-week intracerebroventricular infusion of the antiviral agent ganciclovir (GCV) that kills dividing cells expressing Tk. While GCV drastically reduced the number and growth of hypothalamus-derived neurospheres from adult transgenic mice *in vitro*, it caused hypogonadism *in vivo*. The selective death of dividing tanycytes expressing GFAP indeed caused a marked decrease in testosterone levels and testicular weight, as well as vacuolization of the seminiferous tubules and loss of spermatogenesis. In addition, GCV-treated GFAP-Tk mice showed impaired sexual behavior, but no alteration in food intake or body weight. Our results also show that the selective ablation of GFAP-expressing tanycytes leads to a sharp decrease in the number of gonadotropin-releasing hormone (GnRH)-immunoreactive neurons and blunted LH secretion. Altogether, our data show that GFAP-expressing tanycytes play a central role in the regulation of male reproductive function.

**Main points:** Killing adult hypothalamic GFAP-expressing cells blunts neurosphere formation *in vitro* and leads to GnRH deficiency and hypogonadism *in vivo*. This work pinpoints an unreported role of dividing GFAP-expressing tanycytes in reproductive function.

## INTRODUCTION

In the mammalian brain, neurogenesis consists in the formation of new neurons from neural stem or progenitor cells (NSPCs). In adult, two discrete brain regions defined as neurogenic niches, namely the subventricular zone of the lateral ventricles (SVZ) and the subgranular zone of the hippocampal dentate gyrus (SGZ), have retained this neurogenic potential. Within the specialized microenvironment of these niches, NSPCs exhibit a glial morphology with numerous processes and express astroglial markers such as the glial fibrillary acidic protein (GFAP) (Doetsch, Caille, Lim, Garcia-Verdugo, & Alvarez-Buylla, 1999; Garcia, Doan, Imura, Bush, & Sofroniew, 2004; Imura, Kornblum, & Sofroniew, 2003; Morshead, Garcia, Sofroniew, & van Der Kooy, 2003; Platel, Gordon, Heintz, & Bordey, 2009). *In vitro*, SVZ and SGZ GFAP-positive cells have the ability to generate floating spherical clusters called neurospheres, indicating their stemness potential (Seri, Garcia-Verdugo, McEwen, & Alvarez-Buylla, 2001). Moreover, genetic lineage tracing experiments have confirmed that the progeny of NSPCs derives from cells expressing GFAP (Garcia et al., 2004). One strategy to elucidate the role of the NPSCs is to use the transgenic mouse line GFAP-Tk in which the herpes simplex virus (HSV) thymidine kinase (Tk) is expressed under the control of the Gfap promoter (GFAP-Tk). The antiviral agent ganciclovir (GCV) is a drug which, when phosphorylated by Tk, becomes a toxic metabolite that kills DNA-synthesizing cells. In the GFAP-Tk mouse model, only NSPCs are targeted and killed by GCV, without altering GFAP expression in other cell types such as astrocytes (Garcia et al., 2004). When administered *in vitro* or *in vivo*, GCV therefore selectively eliminates dividing cells expressing both GFAP and Tk, leading to the sharp decrease in SVZ and SGZ neurogenesis (Garcia et al., 2004; Glover, Schoenfeld, Karlsson, Bannerman, & Cameron, 2017; Morshead et al., 2003). Using this strategy, cells expressing GFAP, capable of dividing, were shown to constitute the main population of progenitor cells responsible for constitutive neurogenesis in the two neurogenic niches (Garcia et al., 2004).

Recently, an additional discrete neurogenic niche located in the mediobasal hypothalamus (MBH; for reviews see Yoo & Blackshaw, 2018 and Sharif, Fitzsimons, & Lucassen, 2021) has been documented in various species including human (Pellegrino et al., 2018), mice (Kokoeva, Yin, & Flier, 2007; Kokoeva, Yin, & Flier, 2005), rats (Pencea, Bingaman, Wiegand, & Luskin, 2001; Xu et al., 2005), hamsters (Huang, DeVries, & Bittman, 1998; Kameda, Arai, & Nishimaki, 2003) and sheep (Batailler et al., 2014; Batailler, Derouet, Butruille, & Migaud, 2016; Migaud, Batailler, Pillon, Franceschini, & Malpaux, 2011; Migaud et al., 2010). The hypothalamus, a diencephalic structure located around the 3^rd^ ventricle (3V), is involved in the control of critical physiological functions including food intake and reproduction. Within the hypothalamus, the arcuate nucleus (AN), the location of the central control of appetite and energy balance, contains orexigenic and anorexigenic neurons, including Neuropeptide Y (NPY) and Pro-opiomelanocortin (POMC) neurons, respectively. Besides these metabolic pathways, the hypothalamus also contains GnRH neurons whose cell bodies and nerve endings are located respectively in the hypothalamic preoptic area (POA) and in the median eminence (ME) (Gibson, Ingraham, & Dobrjansky, 2000). GnRH stimulates the production by the pituitary gland of the gonadotrophic hormones, namely the luteinizing hormone (LH) and follicle-stimulating hormone (FSH). In turn, these gonadotrophins regulate gametogenesis and the production of sexual steroids including testosterone and estradiol by the gonads (Knobil, 1990).

In the MBH, the radial glia-like cells lining the ventricular wall, namely the tanycytes, have been identified as the endemic hypothalamic NSPCs that generate new neurons and glial cells in the adult mouse hypothalamus (Lee et al., 2012; Li, Tang, & Cai, 2012; Robins et al., 2013). Tanycytes have their cell body localized in the cellular layer lining the floor of the 3V and send their unique process into the hypothalamic parenchyma. Four subpopulations of tanycytes can be distinguished based on their dorsoventral position, α1, α2, β1 and β2 tanycytes, the more dorsal to the more ventral, down to the ME (Akmayev, Fidelina, Kabolova, Popov, & Schitkova, 1973). All tanycyte subtypes express NSPC markers including the sex-determining region Y-box 2 (Sox2) (Batailler et al., 2014; Lee & Blackshaw, 2012; Li et al., 2012), nestin (Batailler et al., 2014; Wei et al., 2002) and vimentin (Batailler et al., 2014; Bolborea & Dale 2013 ; Mullier, Bouret, Prevot, & Dehouck, 2010), but only α1 and a restricted population of dorsal α2 express the transcript for GFAP and the GFAP protein and have been shown to harbor neural stem cells functional properties (Campbell et al., 2017; Chaker et al., 2016; Robins et al., 2013). In this study, we sought to investigate the role played by dividing GFAP-positive cells in physiological functions controlled by the hypothalamus using the GFAP-Tk transgenic mouse line.

## MATERIALS AND METHODS

### Animals

All the experiments were approved by the Val de Loire animal experimentation ethics committee (CEEAVdL) and were in accordance with the Guidelines of the French Ministry of Agriculture and European regulations on animal experimentation (2010/63/EU). Experiments were performed in accordance with the local animal regulations (authorization N° 2015032613494293 of the French Ministry of Agriculture in accordance with the EEC directive). Experiments were performed on the transgenic mouse strain GFAP-Tk generated as described previously (Bush et al., 1998). Briefly, herpes-simplex virus thymidine kinase (HSV-Tk) is expressed under the control of the mouse *Gfap* promoter. Proliferating GFAP-positive cells expressing the transgene produce toxic nucleotide analogues in the presence of the antiviral agent GCV, which promote their apoptosis. Wild type (WT) and transgenic (Tg) male mice (2 months old) were obtained by mating transgenic females with non-transgenic males. Experiments were performed on four groups, the vehicle-treated WT mice (WT ctr), the GCV-treated WT mice (WT+GCV), the vehicle-treated Tg mice (Tg ctr) and the GCV-treated Tg mice (Tg+GCV).

### RNA extraction and RT-PCR

Total RNA was extracted from the testis and striatum of three 2 months old mice using the RNeasy Mini Kit (Qiagen) and was used for random-primed cDNA synthesis with SuperScript III reverse transcriptase (Thermo Fisher Scientific) following the manufacturer’s instructions. Standard PCR was performed on cDNA aliquots using PlatiniumTaq (Thermo Fisher Scientific) and the specific primers for the thymidine kinase T*k*; Forward primer, CGATGACTTACTGGCGGGTG; Reverse primer, GATACCGCACCGTATTGGCA) and Glyceraldehyde-3-phosphate dehydrogenase (*Gapdh*; Forward primer, CACCATCTTCCAGGAGCGAG; Reverse primer, GTTGAAGTCGCAGGAGACAAC) genes. PCR consisted of a first denaturing step at 94 °C for 2 min, followed by 35 cycles of the following steps at 94 °C for 30 sec, 55 °C for 30 sec, and 72 °C for 45 sec, ending with a 72 °C extension step. PCR products were analysed on a 1.2 % agarose gel containing 0.5 μg/mL ethidium bromide and visualized under a UV transilluminator.

### Neurosphere cultures

Two WT and two Tg male mice (2 months old) *per* experiment (n = 3) were euthanized by cervical dislocation and the region containing the mediobasal hypothalamus was dissected out. Cells were mechanically dissociated using a cell scraper on a nylon membrane and centrifuged 5 min at 1,000 rpm. The resulting single-cell suspensions were plated in a 25 ml flask in DMEM/F12 (Gibco® 21331-020) supplemented with penicillin/streptomycin (PS), B27 and 20 ng/ml each of EGF (Invitrogen 53003-018) and bFGF (Invitrogen 13256-029). Cultures were incubated in humidified chambers with 5% CO_2_ at 37°C. Neurospheres were passaged by centrifuging 5 min at 1,000 rpm and incubating for 7 min in Accutase (Gibco® A11105-01) before mechanically dissociating the spheres into single-cell suspension.

To examine the effect of GCV on neurosphere formation, tertiary neurospheres of WT and Tg mice were cultivated in the presence or absence of GCV (10 µM final concentration, Sigma G2536) for 2 weeks. During this long-term culture, the media was supplemented every two days with growth factors. At the end of the 2 week-culture period, neurospheres were counted and measured for all the conditions.

### *In vivo* GCV administration

Two months old WT ctr (n=9), WT+GCV (n=8), Tg ctr (n=9) and Tg+GCV (n=11) male mice were housed in pairs and the pairs were separated by a Plexiglas partition to maintain visual and olfactory contact. GCV was administrated by intracerebroventricular infusion at a rate of 0.11 µl/hr for 28 days using an osmotic micropump (Model 1004, Alzet®). Micropumps connected to cannula (Brain Infusion Kit 2, Alzet®) were filled with 2 mM GCV or with saline solution and primed for 48 hours at 37°C. For implantation, mice were anesthetized with 100 mg/kg ketamine and 10 mg/kg xylazine and fixed to a stereotaxic frame after loss of reflexes. During surgery, the eyes were protected with ocry-gel and a local anesthetic (procaine) was injected under the skin of the skull. A micropump was introduced subcutaneously and the cannula was implanted into the 3V at 1.7 mm posterior to the Bregma and at a depth of 5 mm (Paxinos atlas). Following surgery, all mice were injected with 0.1 mg/kg morphine analgesic (0.3 mg/ml Buprecare®) and 6 ml/kg injectable antibiotic (10 mg/kg Depocilline).

### Food intake and body weight assessment

Body weights and food intake were measured weekly for 5 weeks starting one week before cannulation at week 0 (W0) until the end of the experiment at week 4 (W4), with week 1 (W1) corresponding to the surgery. To get accurate food intake measurement, mice were housed individually while social interactions (odors, vocalization and sight) were maintained with holes in the cage separator. Food intake was measured by giving a weighted amount of food and measurement of the left over / refusals on a weekly basis (Ali & Kravitz, 2018).

### Behavioral analysis

All the behavioral tests took place in the last week of treatment (W4).

#### Sexual behavior

All sexual behavioral experiments were performed during the dark phase of the dark/light cycle, 1 hour after lights off. Tests were filmed with an infra-red light in a dark room. Two weeks before W0, male mice were placed for a week with a receptive female (in oestrus phase) to gain sexual experience. At W4, males were tested in an open field (50 x 50 cm) with a receptive female for 30 minutes. Females were ovariectomized and implanted with Silastic implants (Dow Corning) containing 50 μg of E2-benzoate (Sigma-Aldrich). Four hours before the tests, females were given a subcutaneous injection of 1 mg of progesterone (Sigma-Aldrich) diluted in 100 μL of oil to induce receptivity (Derouiche, Keller, Duittoz, & Pillon, 2015). The numbers of anogenital investigations, the latencies and frequencies of mounts and intromissions were recorded. The sexual preference of males was also evaluated. Males were placed in the centre of a three-compartment chamber separated by Plexiglas with an opening at the base permitting olfactory and visual contact. Following a 10 min period of habituation, a receptive female (detected by a vaginal smear) and an unfamiliar male were each placed in one of the side compartments of the chamber. The time spent by the tested male mouse near each compartment was recorded for 10 min. Finally, the attractiveness of tested males was measured using the same protocol as above. One oestrus female was placed in the centre of the chamber and a Tg ctr and a Tg+GCV male were placed in the compartments on either side.

#### Anxiety levels

Anxiety level in the four groups was evaluated using the elevated plus maze and the marble burying tests (Meirsman et al., 2016). The elevated plus maze consists of two closed and two open cross-shaped arms (5 cm wide x 40 cm long) elevated 50 cm from the floor. Male mice were placed in the centre of the device and were allowed to explore the arms for 5 minutes. The number of entries and the time spent in each area of the device were recorded, *i*.*e*. the closed arms, the open arms, the distal zones of the open arms and the centre of the device.

In the marble burying test, male mice were placed in a test cage (15 cm wide x 33 cm long) containing fresh bedding. Twenty marbles were distributed evenly in the cage in 5 rows and a lid was placed on top of the cage. Animals were left undisturbed for 15 minutes, after which the number of marbles buried was recorded. A marble was counted as being buried if at least 2/3 of it had been covered by bedding.

### Tissue preparation

At W5, animals were anesthetized and blood was collected from the abdominal artery and serum was frozen for posterior hormone quantification. Following intracardial perfusion with 4% paraformaldehyde, testes, seminal vesicles and preputial glands were dissected out and weighted. Testis samples were fixed by immersion in Bouin’s fixative solution for 48 hours and were embedded in paraffin. Testis blocks were cut in 9 µm thick sections and stained with haematoxylin and eosin for histology. The density of spermatogonia and spermatocytes per seminiferous tubule (number of cells per mm^2^) was assessed. Three images per mouse (2-3 mice per group) and 3 seminiferous tubules per image were used for this quantification.

### Immunohistochemistry

Coronal sections (25 μm thick) cut from the anterior POA in a caudal direction until the premamillary recess, were collected using a cryostat (Leica CM 3050 S). The sections were directly mounted on Superfrost Plus slides (Fisher Scientific, Illkirch, France) and stored at -80°C until used for immunohistochemistry. For all antibodies (Table 1), normal serum IgGs of appropriate species were used as negative controls. For each mouse five sections separated by 100 µm and 160 µm at different rostro-caudal levels of the POA and the MBH respectively were used for immunohistochemistry. To simultaneously permeabilize and block nonspecific binding sites, sections were placed in a solution of 5% normal horse serum and 0.3% Triton X-100 in TBS (TBSTH) for 30 min and incubated in the same buffer containing the primary antibodies (Table 1). They were then incubated with secondary antibodies (Table 1) and mounted in Fluoromount (SouthernBiotech, Birmingham, AL, USA) for observation.

**Table 1.** Primary and secondary antibodies used for immunohistochemistry

| **Primary antibody** | **Manufacturer, species type, cat. no.** | **Unmasking step** | **Dilution** | **Secondary antibody** | **Manufacturer, cat. no.** | **Dilution** |
| --- | --- | --- | --- | --- | --- | --- |
| GFAP | Dako, rabbit polyclonal, #Z0334 |  | 1,1000 | Donkey anti-rabbit IgG, Alexa 488 | Molecular Probes, #A21206 | 1,600 |
| Sox2 | R&D systems, goat polyclonal, #abAF2018 | Sodium borohydure 0.1% | 1,300 | Donkey anti-goat IgG, Alexa 555 | Molecular Probes, #A21432 | 1,600 |
| Vimentin | Millipore, chicken polyclonal, #AB5733 |  | 1,2000 | Donkey anti-chicken IgY, Fluorescein isothiocyanate | Jackson ImmunoResearch #703095155 | 1,600 |
| ERα | Santa Cruz-MC20, rabbit polyclonal, #sc542 | | 1,200 | Donkey anti-rabbit IgG, Alexa 488 | Molecular Probes, #A21206 | 1,600 |
| GnRH | Polyclonal rabbit (no. 19900) |  | 1,10000 | Donkey anti-rabbit IgG, Alexa 488 | Molecular Probes, #A21206 | 1,600 |
| Kisspeptin | Polyclonal rabbit (no. 564) | Citric acid (pH 6) | 1,5000 | Donkey anti-rabbit IgG, Alexa 555 | Molecular Probes, #A31572 | 1,600 |
| NPY | Sigma, rabbit polyclonal, #N9528 | Citric acid (pH 6) | 1,10000 | Donkey anti-rabbit IgG, Alexa 647 | Molecular Probes, #A31573 | 1,600 |
| POMC | Phoenix, rabbit polyclonal, #H02930 | Sodium borohydure 0.1% | 1,400 | Donkey anti-rabbit IgG, Alexa 488 | Molecular Probes, #A21206 | 1,600 |

### Quantification of immunolabelled cells

The quantification of immune-positive cells was performed using a computerized image analysis system, Mercator (Explora Nova, La Rochelle, France), consisting of a microscope (Zeiss Axioscop) equipped with a motorized stage, a fluorescent lamp and a CDD video camera. Under a magnification of 10X, three hypothalamic anatomical regions of interest (ROI), the median eminence (ME), the arcuate nucleus (AN) and the ventromedial hypothalamus (VMH) were determined (Robins et al., 2013). The cell bodies of the GnRH neurons were manually quantified in the POA. Two methods of quantification were used, a manual quantification resulting in a number or a density of immunolabelled nuclei, or an automatic quantification with a grey level threshold to detect the immunolabelled area (Butruille, Batailler, Mazur, Prevot, & Migaud, 2018). The number of GFAP-positive tanycytic cell bodies lining the ventricular wall was quantified in WT ctr and Tg+GCV male mice.

### Image capture for immunohistochemistry

All images (1,024 x 1,024 pixels) for figure preparation were acquired using a confocal microscope (Zeiss, LSM 700, objective 40X) with Zen software (Carl Zeiss, Oberkochen, Germany). Images shown in the figures were pseudo-coloured using LSM Image Browser software (Carl Zeiss, Thornwood, NY), and Photoshop (Adobe Systems, San Jose, CA) was used on the resulting tiff files only to adjust for brightness and contrast.

### Hormone level measurements

Two hours before euthanasia, males were injected intraperitoneally with 15 IU of human chorionic gonadotropin (hCG) (Intervet, France) diluted in physiological serum so that testosterone contained in mouse testis is fully secreted (34). Plasma testosterone concentrations were assayed using a RIA (radioactive immunoassay) using ^3^H-testosterone as previously described (Derouiche et al., 2015). The sensitivity of the assay was 0.06 ng/mL and the intra-assay coefficient of variation was 8.5%.

Plasma cortisol concentrations were measured using a direct radio-immunoassay method as previously described (Orgeur et al., 1999). The sensitivity of the assay was 0.25 ng/mL and the intra-assay coefficient of variation was 8.6%.

Serum FSH levels were measured using a commercial ELISA kit (Endocrine Technologies, Inc., ERKR7014) following manufacturer’s instructions. The sensitivity was 0.05 ng/ml and the intra-assay coefficient of variations was 3.42%.

Serum LH levels were measured using a sensitive sandwich ELISA (Steyn et al., 2013). The sensitivity was 0.25 ng/ml and the intra-assay coefficient of variations was 7.73%.

### Statistical analysis

Statistical analyses were performed with GraphPad Prism5 software (GraphPad Software, San Diego, CA). Data were compared using a non-parametric Kruskal-Wallis test followed by a Dunn’s post-test. A one-way ANOVA followed by a Bonferroni post-test were used to compare the size of the neurospheres after GCV treatment. A two-way ANOVA was used to compare body weights and food intake, and to compare the number of entries and the time spent in each area of the elevated plus maze. A Wilcoxon test was used to compare the sexual preference and the attractiveness of males. Differences were considered statistically significant for a p-value < 0.05. Data presented in the histograms are mean values ± standard error of the mean (S.E.M.).

## RESULTS

### GFAP-expressing cells are the main source of hypothalamic proliferative cells *in vitro*

In the Tg mice model in which the herpes thymidine kinase is expressed under the control of the *Gfap* promoter (GFAP-Tk), double-labelling was used to assess the overlapping expression of GFAP and HSV-Tk at the single-cell level. Confocal analysis of cells showed that 100% of the Tk-positive cells also expressed the astroglial marker GFAP regardless of the rostro-caudal level of the hypothalamus (Fig 1). No Tk-positive/GFAP-negative cells were found and only very few GFAP-positive cells with no detectable level of Tk could be observed. These results confirm the limited expression of Tk in the GFAP-positive cells in the hypothalamus as reported in previous study for the SVZ and SGZ (Garcia et al., 2004).

**Fig 1.**
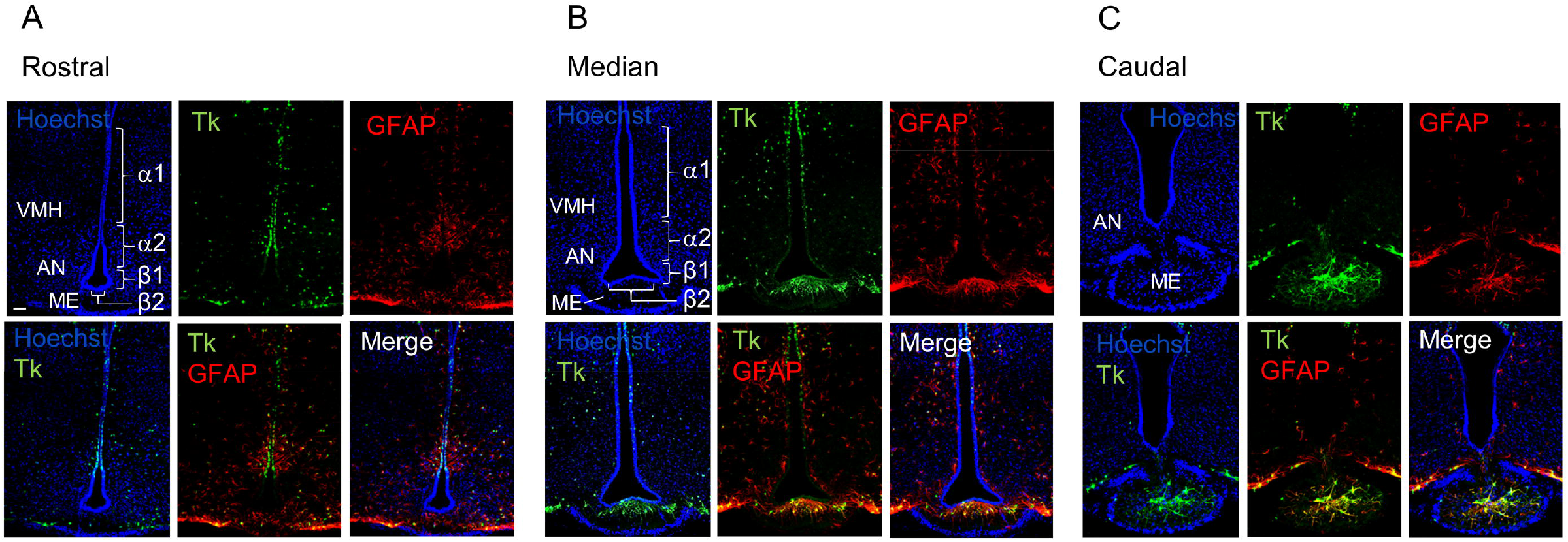
Distribution of GFAP-positive/Tk-positive cells in the MBH of GFAP-Tk male mice. Rostral (A), median (B) and caudal (C) MBH showing for each panel, from left to right and up to down, confocal images of nuclei coloration with Hoechst (blue), Tk (green), GFAP (red), Hoechst/Tk, Hoechst/GFAP expression and the merge image. All the Tk-positive cells express GFAP. They are located in the ependymal layer of the VMH and the dorsal AN, corresponding to the localization of α1 and a subset of α2 tanycytes subtypes. GF AP-positive/Tk-positive cells are also detected in the AN and ME parenchyma. VMH, ventro-medial hypothalamus; AN, arcuate nucleus; ME, median eminence; Tk, thymidine kinase. Scale bar, 20 µm.

Confocal image analysis showed that the majority of Tk-positive/GFAP-positive cells lay in the ependymal layer in the rostral part of the MBH corresponding more specifically to the α-tanycytes subtypes (Fig 1A) and in the MBH parenchyma corresponding to astrocytes (Fig 1A, B and C). A few cells were also observed in astrocytes of the ME (Fig 1B and C). Regarding their anatomical localization, ventricular Tk-positive/GFAP-positive cells will be referred to as GFAP-positive tanycytes in the remainder of this article.

To determine the relative contribution of GFAP-expressing cells to NSPCs isolated from hypothalamic tissue, we used the neurosphere assay, which constitutes the best *in vitro* assay for detecting the presence of putative neural progenitor cells (Marshall, Reynolds, & Laywell, 2007). The number of neurospheres was comparable in non-transgenic mice (WT ctr), in GCV-treated non-transgenic (WT+GCV) mice or in saline-treated transgenic mice (Tg ctr; Fig 2A, B and C respectively). In contrast, GCV almost completely prevented neurosphere formation from adult hypothalamic tissue derived from transgenic mice (Tg+GCV; Fig 2D). The GCV treatment reduced the number of neurospheres from Tg+GCV mice by 89.2±9.5% (Fig 2E). Furthermore, neurospheres produced from Tg male mice treated with GCV were significantly smaller than those from control mice (Fig 2F; Bonferroni multiple comparison, p<0.0001). These results show that the ability of adult hypothalamic tissue to produce neurospheres *in vitro* was drastically reduced following transgenic-targeted elimination of dividing GFAP-positive cells.

**Fig 2.**
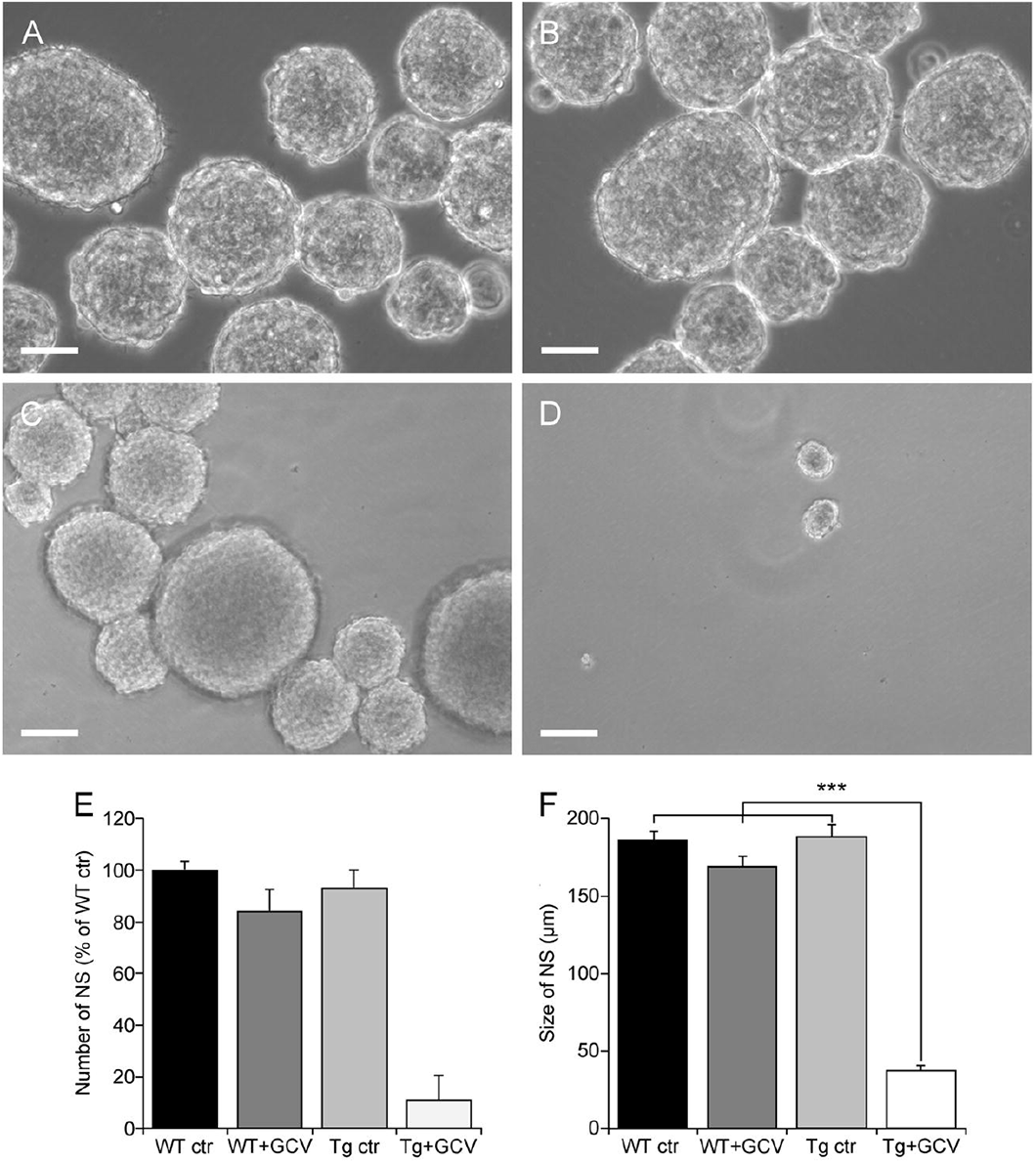
GFAP-expressing cells are the predominant source of proliferative cells within the hypothalamus. Representative images of floating hypothalamic tertiary neurospheres (NS) derived from wild type mice (WT ctr; A), wild type mice receiving a ganciclovir treatment (WT+GCV; B) GFAP-Tk Tg mice (Tg ctr; C) and GFAP-Tk Tg mice treated with an i.c.v. ganciclovir injection (Tg+ GCV; D). Scale bar, 100 µm. Number (E) and size (F) of hypothalamic neurospheres after a 2-week culture period in presence of GCV. Data are expressed as the mean±SEM, WT n=2 and Tg mice n>3 for each group, n=3 experiments. ***p<0.001.

### *In vivo* ablation of GFAP-positive tanycytes modified the expression of NSPC markers in the MBH

To determine whether GFAP-expressing tanycytes could constitute the major pool of NSPCs in the adult hypothalamus, two months old WT and Tg mice were subjected to 4-week saline and GCV infusion in the 3V using stereotaxically implanted cannulas and osmotic minipumps (WT ctr; WT+GCV; Tg ctr; Tg+GCV).

In order to assess the effect of the GCV treatment on the expression of NSPC markers, brain slices were cut and immunohistochemical analyses were performed. The expression of NSPC markers, including GFAP (Fig 3A-D), vimentin (Fig 3E-G) and Sox2 (Fig 3H-K) was quantified using immunolabelling techniques.

**Fig 3.**
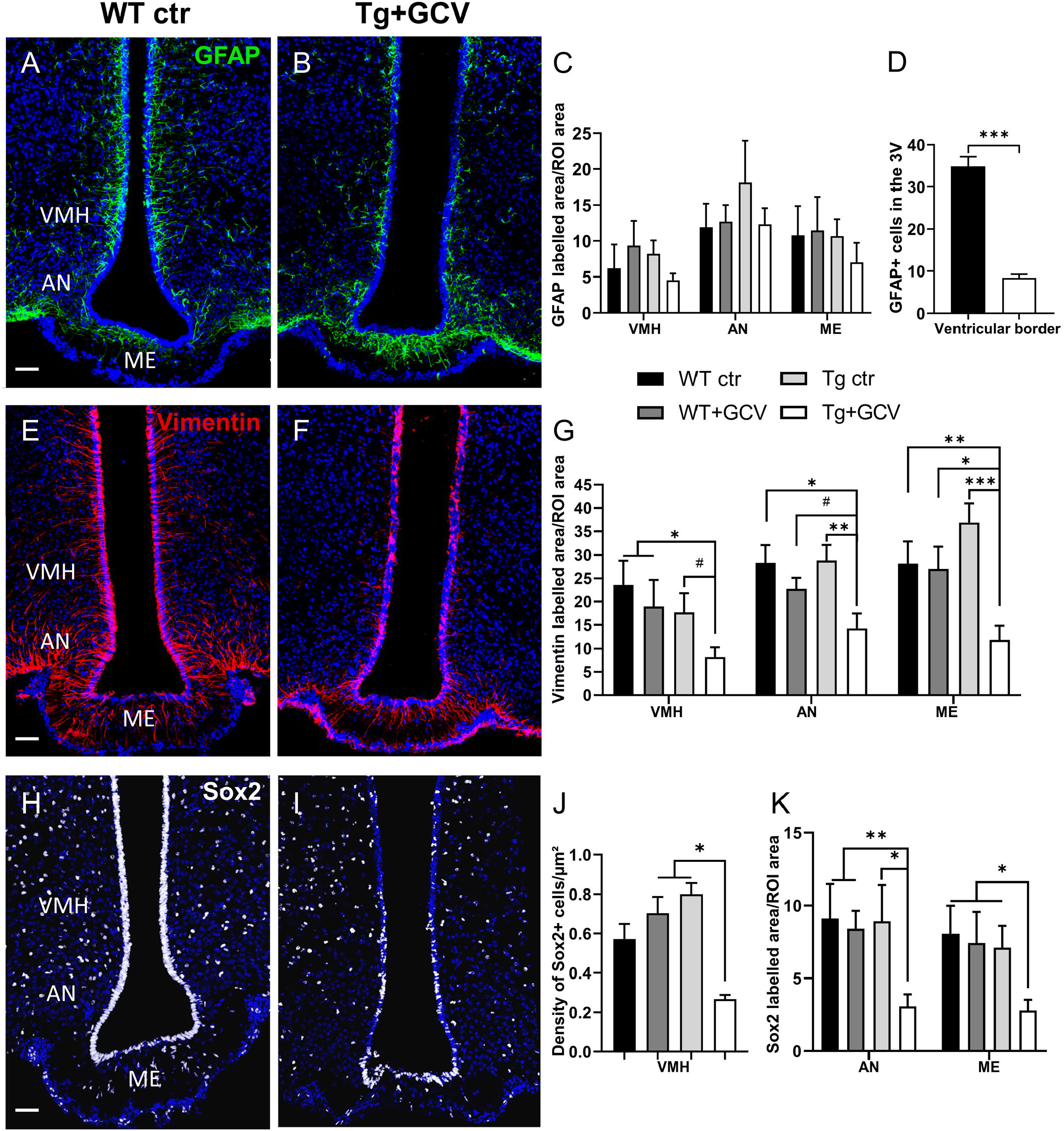
*In vivo* ablation of GFAP-expressing tanycytes impaired expression of NSPCs markers within the MBH. Representative images of GFAP expression measured in the mediobasal hypothalamus of control (A) and Tg+GCV male mice (B). Statistical analysis revealed no variation of GFAP expression in the VMH, AN and ME (C). In contrast, the number of GFAP-positive tanycytes located in the ventricular border is statistically lower in Tg+GCV male mice as compared to control mice (D). Confocal images of Vimentin expression in the mediobasal hypothalamus of control (E) and Tg+GCV male mice (F). Tg+GCV male mice display a significantly decreased Vimentin expression in the VMH, NA and ME (G). Representative images of Sox2 expression in the mediobasal hypothalamus of control (H) and Tg+GCV male mice (I). Statistical analysis revealed significantly less Sox2 positive cells in the ventromedial hypothalamus (VMH) (J), in the AN and in the ME (K) in Tg+GCV male mice. Data are expressed as the mean±SEM, n≥5 for each group. *p<0.05, **p<0.01. ***p<0.001, #p=0.06 or 0.07. Scale bars, 50 µm.

In Tg mice, the GCV treatment had no effect on the labelling of GFAP in the parenchyma of the VMH (Kruskal-Wallis, p>0.05), AN (Kruskal-Wallis, p>0.05) or ME (Kruskal-Wallis, p>0.05) globally, when compared with the WT, WT+GCV and Tg ctr groups (Fig 3C). However, analysis of the GFAP-positive labelling located in the wall of the 3V revealed that the number of GFAP-positive tanycytic cell bodies is dramatically decreased in Tg+GCV mice when compared to WT ctr littermates (Mann-Whitney, p=0.0002; Fig 3D). These results thus suggest that GCV treatment *i*.*c*.*v*. selectively promotes the ablation of GFAP-positive tanycytes along the wall of the 3V without altering the GFAP-positive astrocytes of the parenchyma. The expression of vimentin (Fig 3E-F), a type III intermediate filament protein expressed in neural stem cells, was significantly reduced in the VMH, AN and ME of the Tg+GCV group as compared to the WT, WT+GCV and Tg ctr groups (VMH: Kruskal-Wallis, p=0.05; Dunn’s, p<0.05; AN: Kruskal-Wallis, p=0.03; Dunn’s, 0.008<p<0.07; ME: Kruskal-Wallis, p=0.003; Dunn’s, 0.0004<p<0.01 Fig 3E-G). These results intriguingly suggest that the ablation of dividing GFAP-positive tanycytes does not only cause depletion in the α-tanycyte population in the wall of the 3V, but also in the β-tanycytes, which form the floor of the 3V in the ME. In addition to a marked depletion of Sox2-positive tanycytes (Fig 3H, I), the density of Sox2-positive cells in the parenchyma of the VMH (Fig 3J), was significantly lower in the Tg+GCV group than in the WT+GCV and Tg ctr groups (Kruskal-Wallis, p=0.01; Dunn’s, p=0.04 and p=0.05 respectively Fig 3J). A significant decrease in Sox2-positive cells density was also observed in the AN (Kruskal-Wallis, p=0.02; Dunn’s, 0.005< p<0.03) and the ME (Kruskal-Wallis, p=0.04; Dunn’s, 0.02<p<0.04; Fig 3K). Altogether, these results raise the possibility that GFAP-expressing tanycytes may play a role in the renewal of all tanycytic populations in the ventricular wall, as well as of transient amplifying cells in the parenchyma, in the adult tuberal region of the hypothalamus.

### *In vivo* ablation of hypothalamic GFAP-positive tanycytes did not affect body weight and food intake and has no effect on anorexigenic/orexigenic peptide expression

As adult hypothalamic neurogenesis has been associated with the control of food intake and metabolism (Kokoeva et al., 2005; Lee et al., 2012; Li, Tang, Purkayastha, Yan, & Cai, 2014; Pierce & Xu, 2010), animals and their food intake were therefore weighed daily from week 0 (W0) to week 4 (W4) after GFAP-positive tanycytes ablation. A two-way ANOVA demonstrated that the GCV administration did not modify the body weight or food intake of the Tg-GCV mice when compared to the three control groups of mice (p>0.05, Fig S1A, and p>0.05, Fig S1B, respectively).

The balance between anorexigenic neurons expressing proopiomelanocortin (POMC) and orexigenic neurons expressing neuropeptide Y (NPY) mainly mediates metabolic activity including feeding and body weight regulation. In transgenic GCV-treated mice, the number of neurons immunolabelled for POMC (WT ctr: 37.78±8.45; WT+GCV: 32.06±9.31; Tg ctr: 45.84±10.9; Tg+GCV: 30.91±8.08; Kruskal-Wallis, p>0.05; Fig S1C) and NPY (WT ctr: 10.45±1.81; WT+GCV: 8.97±0.43; Tg ctr: 13.96±1.3; Tg+GCV: 13.72±2.43; Kruskal-Wallis, p>0.05; Fig S1D) did not differ from the control groups. Taken together these data indicate that ablation of hypothalamic GFAP-positive tanycytes did not cause any marked alteration in the regulation of body weight or food intake during the four weeks of treatment. However, we cannot exclude metabolic consequences at a longer term, as previously demonstrated by the decrease in POMC neurons and the alteration of body weight and food intake 3 months and 10 months respectively after induced inflammation in hypothalamic NSCs (Li et al., 2012).

### Ablation of hypothalamic GFAP-positive tanycytes causes hypogonadism

We next investigated the role of adult GFAP-expressing tanycytes on reproduction, a major neuroendocrine function controlled by the hypothalamus. The effect of 4-week GCV treatment on the weight of testes and seminal and preputial glands was examined. Tg+GCV mice showed a stricking decrease in testis weight (Kruskal-Wallis, p<0.0001; Dunn’s, p<0.05; Fig 4A). Following a human chorionic gonadotropin (hCG) injection, a stimulation that elicits the release of the total testosterone content from the testis (Derouiche et al., 2015), a significant decrease of about 50% of the mean plasma testosterone levels was observed in Tg+GCV compared to control groups (Kruskal-Wallis, p=0.04, Dunn’s, p<0.05; Fig 4B). Anatomical analysis of the testis sections showed that the three control groups, *i*.*e*. WT ctr (Fig 4C), WT+GCV (Fig 4D) and Tg ctr (Fig 4E), had normal seminiferous tubules. In contrast, the GCV treatment of Tg mice resulted in severe morphological alterations including vacuolization of seminiferous epithelium and no visible spermatozoa in the tube lumen (Fig 4F). A morphological analysis of the seminiferous tubules of the Tg+GCV mice showed that they only contained Sertoli cells, spermatogonia and some spermatocytes. The stages downstream the spermatogonia stage were affected by the GCV treatment and no spermatids or spermatozoa could be observed in the tubules suggesting that in male Tg+GCV mice, spermatogenesis had stopped at the leptotene spermatocyte stage which is known to be highly sensitive to testosterone variations (Chang et al., 2004). Some of the seminiferous tubules also contained apoptotic vesicles corresponding to phagocytosis of spermatogonia by the Sertoli cells. The quantification of the different cell types in the seminiferous tubes uncovered that the Tg+GCV mice have significantly less spermatogonia and spermatocytes than the control mice (Kruskal-Wallis, p=0.0002; Dunn’s, p<0.05; Fig S2). Moreover, no *Gfap* expression was detected in the testis of the control mice (data not shown) and the expression of the *tk* gene was undetectable in testis of Tg mice as shown by RT-PCR (Fig S3), indicating that anatomical defects observed in the testis were not due to a direct peripheral effect of the ganciclovir. These findings demonstrate that the ablation of GFAP-positive tanycytes in Tg+GCV male mice causes rapid and profound disruption of spermatogenesis and severe degradation of the testicular morphology.

**Fig 4.**
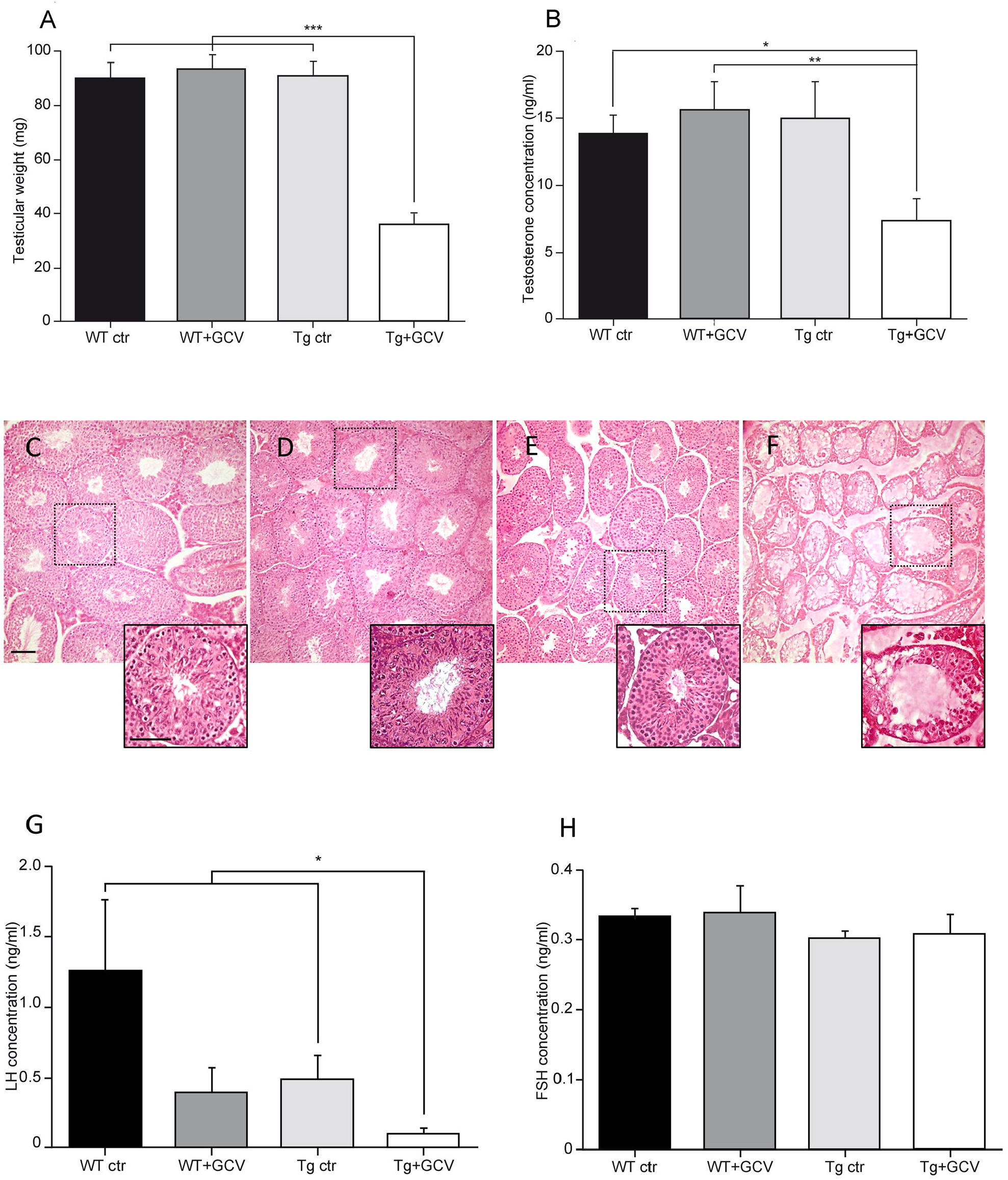
Alteration of gonadal functions following ablation of GFAP-positive tanycytes. Tg+GCV male mice display decreased testicular weight (A) and plasma testosterone concentrations after hCG (human chorionic gonadotropin) injection (B). Testicular histology of WT ctr (C), WT +GCV (D), Tg ctr (E) and Tg+GCV (F) male mice revealed alterations and vacuolization of seminiferous tubules after the ablation of GFAP positive NSPCs, scale bars, 50 µm Insets show a higher magnification image of a single seminiferous tube for each group of mice. Scale bar, 200 µm. Tg+GCV male mice display decreased serum LH concentration (G) while serum FSH concentration is not affected by the treatment (H). Data are expressed as the mean±SEM, n≥3 for each group. *p<0.05, **p<0.01, ***p<0.001.

In addition, Tg+GCV mice showed a decrease in the weight of preputial glands which produce pheromones (WT ctr: 62.67±4.28; WT+GCV: 68.34±5.35; Tg ctr: 61.53±3.13; Tg+GCV: 47.83±3.39; Kruskal-Wallis, p=0.03; Dunn’s, p<0.05), whereas no change in the seminal gland weight was observed between the four groups (WT ctr: 227.2±15.17; WT+GCV: 226.4±13.56; Tg ctr: 254.3±32.02; Tg+GCV: 219.32±4.5; Kruskal-Wallis, p>0.05).

The release of testosterone is triggered by the action of the pituitary hormone LH on Leydig cells, we therefore analysed circulating LH levels. In Tg+GCV male mice, a significant decrease in the mean serum LH concentrations was observed (Kruskal-Wallis, p=0.04; Dunn’s, p<0.05; Fig 4G) compared to control groups except for WT+GCV mice. In contrast, no significant difference was found in the mean levels of FSH, which regulate Sertoli cell function (Kruskal-Wallis, p>0.05; Fig 4H) in the male mice of the four groups. Together these data strongly suggest that 4-week delivery of GCV in the third ventricle of GFAP-Tk mice causes severe hypogonadotropic hypogonadism.

We next sought to investigate the consequences of tanycytic depletion on the function of the GnRH neuronal network within the hypothalamus. Tanycytes of the ME tightly control the access of GnRH nerve terminals to the pituitary portal blood vessels (Parkash et al., 2015; Prevot et al., 1999). Furthermore, tanycytes in the wall of the 3V have recently been shown to be able to modulate the activity of neurons in the AN (Bolborea, Pollatzek, Benford, Sotelo-Hitschfeld, & Dale, 2020), which is a key site for the feedback action of gonadal steroid in the hypothalamus where reside kisspeptin neurons. While neither Kisspeptin immunoreactivity nor the number of estradiol receptor alpha (ERα)-positive cells were affected by GCV treatment in the AN (Kruskal-Wallis, p>0.05; Fig S4A, B), density of GnRH neuronal fibers were found to be significantly decreased in the ME of GCV-treated mice (Kruskal-Wallis, p=0.02, Dunn’s, p<0.05; Fig 5A-C). Surprisingly, this decreased density of GnRH neuronal fibers in the ME was associated with a 50% loss in the number of GnRH-immunoreactive neuronal cell bodies in the preoptic region (Kruskal-Wallis, p=0.002, Dunn’s, p<0.05; Fig 5D-F). These results intriguingly suggest that morphofunctional interaction between GnRH axon terminals and tanycytes is not only required for the control of GnRH release into the pituitary portal blood but also GnRH expression and/or GnRH neuronal survival in the hypothalamus.

**Fig 5.**
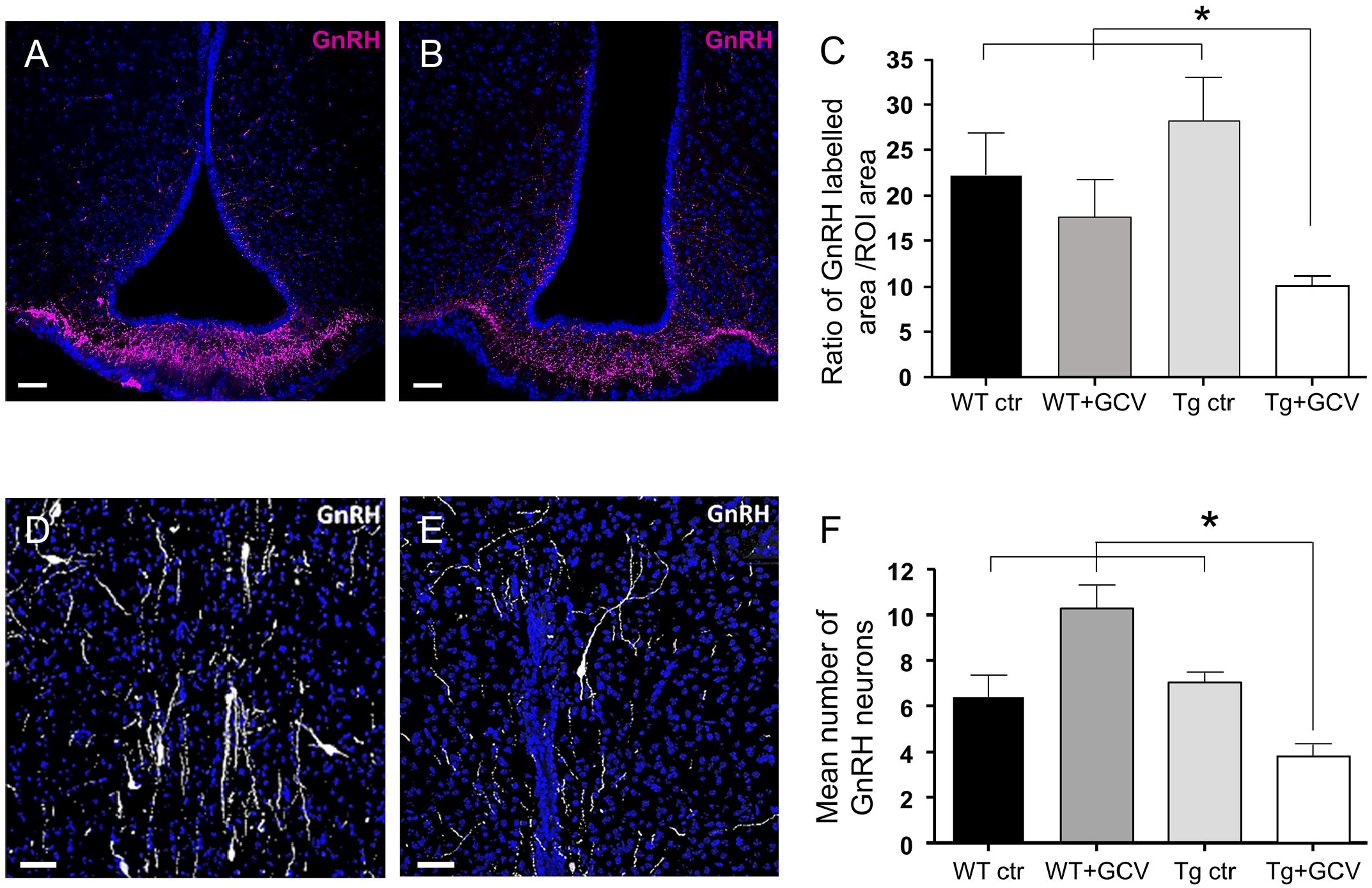
Effect of the ablation of GFAP-positive tanycytes on GnRH immunoreactivity in the preoptic region and the median eminence. (A-B) Representative images of GnRH fibers in the median eminence of control (A) and Tg+GCV (B) male mice. Statistical analysis revealed a significantly decreased GnRH expression in Tg+GCV mice as compared to the three control groups (C). (D-E) Representative images of GnRH neurons in the preoptic area (POA) of control (D) and Tg+GCV (E) male mice. (F) Mean number of GnRH neurons in the POA per slices, n≥3 slides per group. Statistical analysis revealed a significantly decreased GnRH neuron number in the POA in Tg+GCV male mice (F). Data are expressed as the mean±SEM. *p<0.05. Scale bars, 20 µm.

### GFAP-expressing tanycytic depletion alters male sexual behaviour

To further explore the consequences of tanycytic depletion on reproduction, three main well-established paradigms of sexual behaviour in male mice, namely the frequency of anogenital investigations, the latency to the first mount and intromission and the frequency of intromissions were analysed. To achieve this, male mice of the four experimental groups were exposed to receptive females for 30 minutes. In this test, the GCV treatment did not induce any change in the frequency of anogenital investigation in transgenic male mice (Kruskal-Wallis, p>0.05; Fig 6A). In contrast, the Tg+GCV male mice exhibited increased latencies to the first mount (Kruskal-Wallis, p=0.01; Dunn’s, p<0.05; Fig 6B) as well as for the first intromission (Kruskal-Wallis, p=0.03 Dunn’s, p<0.05; Fig 6C) together with a lower frequency of intromissions (Kruskal-Wallis, p=0.008; Mann-Whitney, p<0.05; Fig 6D). The attractiveness of Tg ctr and Tg+GCV males towards receptive females was also analysed and showed that receptive females spent as much time with Tg ctr males as with Tg+GCV males (Wilcoxon, p>0.05; Fig 6E), suggesting that Tg+GCV males were as attractive for females as control males. Conversely, to test whether the decrease in sexual performance of the Tg+GCV male mice was due to a lack of sexual attraction towards females, we performed a sexual preference test by exposing male mice of the four groups to both an unfamiliar male and a receptive female. Males from all groups spent more time near the female than near the unfamiliar male (Wilcoxon, WT ctr p=0.05, WT+GCV p=0.03, Tg ctr p=0.001 and Tg+GCV p=0.004; Fig 6F), indicating that GCV treatment did not alter the sexual preference of the treated males for females. Altogether these results show that GCV-treatment leads to an impairment of the copulative behaviour without affecting sexual preference and attractivity in male mice.

**Fig 6.**
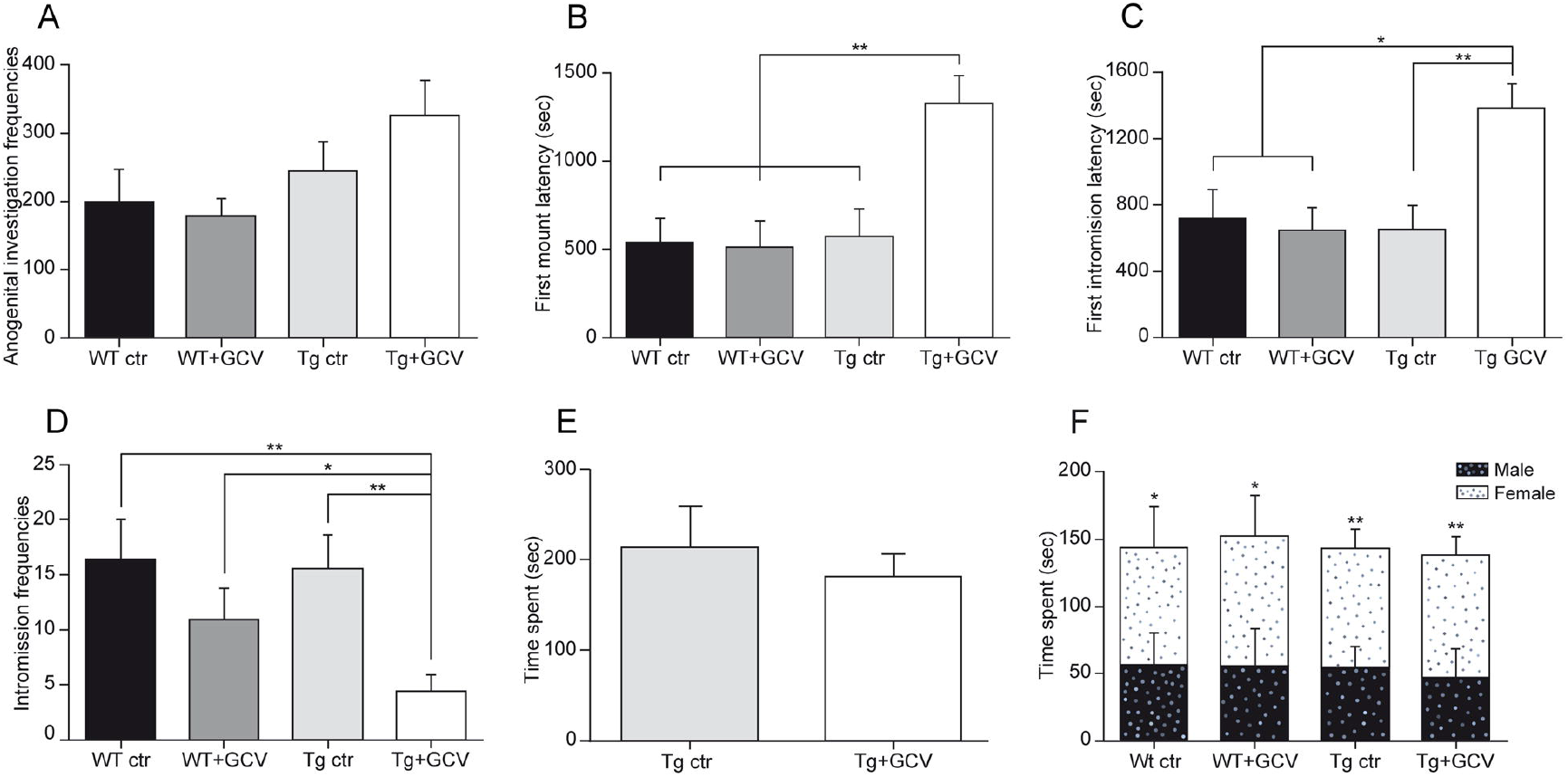
Ablation of GFAP-expressing tanycytes affected male sexual behaviour. The number of anogenital investigations is not altered by the ablation of GFAP-expressing tanycytes (A). The latencies to first mount (B) and to first intromission (C) are increased in Tg+GCV male mice, while the intromission frequencies (D) are decreased after ablation of GFAP tanycytes. Receptive females spend as much time in close contact to either Tg ctr or Tg+GCV male mice (E). Males of the four groups spend more time in close contact to a receptive female than an unfamiliar male (F). Data are expressed as the mean±SEM, n≥6 for each group. *p<0.05, **p<0.01.

### Ablation of GFAP-positive tanycytes does not alter anxiety levels

The hypothalamus regulates stress and anxiety responses via the corticotropic axis and tanycytes are also known to be morphologically associated with corticotropin releasing hormone (CRH) axon terminals (see for review Prevot et al., 2018). To test whether ablating GFAP-positive tanycytes might also lead to depressive/anxiety behaviour, two behavioural studies were conducted, namely the elevated plus maze and the marble burying tests. The GCV treatment did not modify the number of entries into each zone of the elevated plus maze (two-way ANOVA, p>0.05; Fig S5A) and Tg+GCV male mice spent as much time in each area of the elevated plus maze device as the three control groups (two-way ANOVA, p>0.05; Fig S5B). The marble burying test, an anxiety test that partly depends on hippocampal function (Deacon, 2006) showed that animals buried the same number of marbles whichever group they belonged to (Kruskal-Wallis, p>0.05; Fig S5C). Additionally, no significant difference was detected in the mean plasma cortisol concentrations between the four groups (Kruskal-Wallis, p>0.05; Fig S5D), consistent with the findings that the GCV treatment had no effect on the anxiety level in transgenic mice.

## DISCUSSION

It is now well established that adult dividing GFAP-positive cells constitute the unique pool of NSPCs in the SVZ and SGZ (Doetsch et al., 1999; Garcia et al., 2004; Imura et al., 2003; Morshead et al., 2003; Platel et al., 2009; Seri et al., 2001) and their ablation results in a depletion of adult neurogenesis within the two canonical niches (Garcia et al., 2004; Morshead et al., 2003). In the hypothalamus, tanycytes have been described as the NSPCs capable of generating neurons and glial cells (Sousa-Ferreira, de Almeida, & Cavadas, 2014; Yoo & Blackshaw, 2018). Despite strong morphological similarities, two recent single-cell RNA sequencing (scRNA-Seq) studies showed much greater molecular diversity than previously thought among tanycytes (Campbell et al., 2017; Chen, Wu, Jiang, & Zhang, 2017). Although all populations express Sox2, Vimentin and Nestin, only the α-tanycyte subtypes express GFAP (Campbell et al., 2017; Robins et al., 2013), while Fgf10 (fibroblast growth factor 10) and BLBP (Brain Lipid-Binding Protein) are only expressed by the β subtypes (Goodman et al., 2020; Haan et al., 2013; Robins et al., 2013), suggesting distinct roles of these two NSPCs populations. In the current study, we used a Tk-transgenic mouse line in which only dividing GFAP-positive NSPCs are selectively removed while non-stem GFAP-positive astrocytes are spared by ganciclovir treatment (Garcia et al., 2004).

In the hypothalamus, dividing GFAP-positive NSPCs that were selectively targeted in our study correspond mainly to one of the mitogenic regions identified in the tanycytic layer lining the 3V wall, namely the α1 plus a dorsal subset of α2 tanycytes (Robins et al., 2013). *In vitro*, selective ablation of dividing GFAP-expressing cells induced a strong reduction in the neurospherogenic capacities of this region as measured in the neurosphere assay by a sharp reduction in both the number and the size of neurospheres. *In vivo*, ablation of dividing GFAP-expressing tanycytes resulted in decreased expression of NSPC markers including vimentin and Sox2 thoughout the MBH of Tg mice. In agreement with these results, α-tanycytes expressing GFAP and Sox2 were shown to self-renew, give rise to β-tanycytes and generate neurons and astrocytes (Robins et al., 2013; Rodriguez et al., 2005; Xu et al., 2005). These results support the assumption that GFAP-expressing α-tanycytes proliferate and show strong neurospherogenic capacities *in vitro* (Robins et al., 2013; Rodriguez et al., 2005; Xu et al., 2005), thus behaving like NSPCs, similarly to those of the two canonical niches, the SVZ and the SGZ (Garcia et al., 2004).

Several studies have reported that newborn cells, most likely generated from tanycytes, integrate the hypothalamic neuronal networks of the AN and ME and are engaged in the regulation of energy homeostasis (Haan et al., 2013; Kokoeva et al., 2005; Lee et al., 2012; Li et al., 2012). A decrease in hypothalamic new cell generation induces obesity and the development of metabolic disorders associated with insulin resistance (Li et al., 2012). In turn, obesity induced by a high fat diet was shown to impair hypothalamic neurogenesis and disrupt energy balance (Li et al., 2012; Sousa-Ferreira et al., 2014). In addition, a 2-month high fat diet induces a significant increase in the proliferation of β-tanycytes and the production of new cells in the ME (Lee et al., 2012). In order to determine whether hypothalamic GFAP-expressing tanycytes are implicated in metabolism regulation, we explored the effect of their ablation on the regulation of body weight and food intake in transgenic male mice. The current study demonstrates that the suppression of GFAP-expressing tanycytes had no effect on both parameters, nor on the number of anorexigenic and orexigenic neurons *i*.*e*. POMC and NPY neurons respectively. Two hypotheses could be formulated to explain these results. First, this targeted population of NSPCs, corresponding mainly to the α1 and dorsal α2 tanycytes, would not be directly involved in the regulation of food intake, unlike the β tanycytes (Lee et al., 2012; Li et al., 2012). Alternatively, no effect was detected because the animals were unchallenged. The exploration of whether a diet challenge could affect food intake in GFAP-expressing tanycyte-depleted mice might bring insight to this question.

We then investigated the involvement of hypothalamic GFAP-expressing tanycytes in the control of reproduction, a function also orchestrated by the hypothalamus. In a recent study in sheep, a seasonal mammal, we showed that administration of the antimitotic drug AraC in the 3V downregulates the production of hypothalamic new neurons identified by the expression of Doublecortin and modifies the timing of reproduction (Batailler et al., 2018). In a model of aged mice, GnRH was shown to stimulate the hypothalamic mitogenic activity (Zhang et al., 2013). These data suggest a strong link between the hypothalamic neural stem cell niche and the neural circuits controlling reproduction. The current study shows that the sexual behaviour and sexual performance of male Tg+GCV mice are strongly altered after 4-week of GCV treatment and thus without affecting the neural circuits involved in sexual partner olfaction and recognition. This phenotype does not result from a higher level of anxiety in Tg+GCV mice as they exhibit plasma cortisol concentrations and levels of anxiety comparable to those in the control groups.

A thorough analysis of the reproductive system of the Tg+GCV male mice showed a significant decrease in testis weight and a drastic reduction in testosterone secretion by the Leydig’s cells. In turn, the drop of testosterone secretion triggered the vacuolization of the seminiferous tubules likely due to the cessation of spermatogenesis and severe hypogonadism. These decreased testosterone levels are also likely the cause of the strong alteration in sexual behaviour seen in GCV-treated transgenic males.

Testosterone inhibits GnRH/LH secretion via a negative feedback onto the hypothalamus. The drop in testosterone levels being associated with decreased circulating LH levels in GCV-treated mice strongly suggests that the hypogonadism in these mice is due to a central defect. In agreement, we found that GCV-mediated depletion in tanycytes results in GnRH deficiency. GnRH immunoreactivity was indeed seen to be markedly decreased both at the GnRH neuronal cell bodies in the POA and the termination field of GnRH neurons in the ME. Interestingly, GCV treatment in transgenic mice did not affect FSH levels. These results are consistent with previous studies in which the use of GnRH antagonists in humans induce an immediate decrease in LH and testosterone secretions, while the decrease in FSH is delayed and weaker because of its short half-life (Hall et al., 1988; Pavlou et al., 1986). Interestingly, these antagonist treatments cause a rapid hypogonadism phenotype that is dependent on fluctuations in pituitary hormones and testosterone, comparable to our data (Hall et al., 1988; Pavlou et al., 1986). In ewes (Lincoln & Fraser, 1979), non-human primates (Pineda et al., 1983), and ovariectomized rats (Culler & Negro-Vilar, 1986, 1987), the administration of a GnRH antagonist as well as the hypothalamic-pituitary disconnection (Hamernik, Crowder, Nilson, & Nett, 1986) suppressed pulsatile LH secretion but had a minimal effect on FSH secretion, indicating that GnRH differentially controls the secretion of LH and FSH. However, a GCV treatment of more than 4 weeks of Tg mice might also have altered FSH secretion. Together with recent findings showing that the administration of the antimitotic AraC in the lateral ventricle of female rats alters the neural circuitry that controls adult GnRH/LH release and reduces the preovulatory surge in LH release (Mohr, DonCarlos, & Sisk, 2017), our results point to a role for the adult hypothalamic neural stem cell niche in the control of the reproductive function.

We do not know the cellular mechanisms inducing GnRH deficiency following the suppression of hypothalamic GFAP-expressing tanycytes. Kisspeptin-54 being the most potent secretagogue of the GnRH system (Messager et al., 2005), and having been reported to induce a reduction in testicular weight as well as testicular degeneration in adult male rats when chronically administrated (Thompson et al., 2006), was one of the candidates. However, Kisspeptin expression in fibers located in the AN did not appear to be altered in GCV-treated transgenic mice. Similarly, expression of gonadal steroid receptors such as the estrogen receptor-α (ERα) appeared to be unaffected. Alternately, a more direct hypothesis involves the β-tanycytes located in the ME, which mediate the release of GnRH into the pituitary portal blood vessels by controlling the direct access of GnRH nerve endings to the vascular wall (see for review Prevot et al., 2018) but also by setting up specific communication channels with GnRH nerve terminals (see for review Clasadonte & Prevot, 2018) and supporting neuronal survival (Chauvet, Privat, & Alonso, 1996; Prieto, Chauvet, & Alonso, 2000). However, this view is somewhat contradicted by a recent study reporting the physiological consequences of selective β-tanycytes ablation (Yoo et al., 2020). Although it shows, as in the present study, that β-tanycytes ablation has no effect on food intake and body weight, the authors did not observe any significant changes in serum levels of LH and FSH. This discrepancy could be explained by the fact that the animals in this study were fed a diet containing tamoxifen for 3-weeks; tamoxifen being a potent modulator of gonadal steroid receptors leading to confounding effects on the activity of the gonadotropic axis. In particular, tamoxifen has been clinically shown to increase androgen levels and sperm concentration in males with idiopathic oligozoospermia (Dimakopoulou, Foran, Jayasena, & Minhas, 2020).

GFAP-expressing α1-and dorsal α2-tanycytes may communicate with GnRH fibers traveling towards the ME, but do not interact morphologically with GnRH nerve terminals in the ME. However, their ablation caused a marked alteration in β-tanycyte number and/or morphology as evidenced by the significant decrease in vimentin labelling 4 weeks after GCV treatment (Fig 3E-G). These results are in agreement with the finding that α-tanycytes expressing GFAP self-renew and give rise to β-tanycytes (Dimakopoulou et al., 2020; Robins et al., 2013; Rodriguez et al., 2005; Xu et al., 2005). The immunohistological changes observed in our study likely resulting in significant structural alterations may hamper the tanycyte-to-GnRH-neuron communication processes controlling reproduction (Prevot et al., 2018). Hence, one is tempted to speculate that this alteration in the structure and function of hypothalamic tanycytes may also account, at least in part, for the loss of GnRH immunoreactivity in the ME with possible consequences for overall expression of GnRH in the cell bodies or for the very survival of GnRH neurons in the preoptic region. This phenomenon, which could be due to an alteration of the ability of tanycytes to fight against systemic inflammation at the blood-cerebrospinal barriers (Bottcher et al., 2020; Li et al., 2012; Zhang et al., 2013), affecting fitness and, maybe, viability of neuroendocrine systems (Osterstock et al., 2014) or promoting changes in gene-miRNA micronetworks (Messina et al., 2016; Zhang et al., 2017) remains to be explored.

In conclusion, our results established the importance of α-tanycytes in the production and maintenance of the NSPC pool of the entire MBH. In particular, they show that the elimination of these GFAP-expressing tanycytes severely impairs the activity and function of GnRH neurons leading to hypogonadotropic hypogonadism and alters sexual behaviours in male mice, highlighting their key role in the control of reproduction.

## Acknowledgments

L. Butruille received a grant from the Région Centre. This work was funded by the PHASE department of INRAE. This project was funded by the Agence Nationale de la Recherche ANR-16-CE37-0006 (to M.M) and the European Research Council (ERC) Synergy Grant WATCH No 810331 (to VP). The authors thank the PAO experimental unit No. 1297 (EU0028; INRAE Val de Loire) for animal care, Anne-Lyse Lainé and the members of the hormonal assay platform, Nouzilly, France, for conducting testosterone and cortisol assays and the PIC platform INRAE-Val de Loire UMR PRC. The authors wish to thank Dr Alexandre Surget, Dr Matthieu Keller, Dr Julie Le Merrer and Dr Jérôme Becker for technical and scientific advice and for their help with behavioural testing and Professor Michael Sofroniew for kindly donating the Tk antibody.

## REFERENCES

Akmayev, I. G., Fidelina, O. V., Kabolova, Z. A., Popov, A. P., & Schitkova, T. A. (1973). Morphological aspects of the hypothalamic-hypophyseal system. IV. Medial basal hypothalamus. An experimental morphological study. Z Zellforsch Mikrosk Anat, 137(4), 493–512. doi: 10.1007/BF00307226

Ali, M. A., & Kravitz, A. V. (2018). Challenges in quantifying food intake in rodents. Brain Res, 1693(Pt B), 188–191. doi: 10.1016/j.brainres.2018.02.040

Batailler M, Droguerre M, Baroncini M, Fontaine C, Prevot V, & Migaud M. (2014). DCX-expressing cells in the vicinity of the hypothalamic neurogenic niche: a comparative study between mouse, sheep, and human tissues. J Comp Neurol, 522(8), 1966–1985.

Batailler, M., Chesneau, D., Derouet, L., Butruille, L., Segura, S., Cognie, J., … Migaud, M. (2018). Pineal-dependent increase of hypothalamic neurogenesis contributes to the timing of seasonal reproduction in sheep. Sci Rep, 8(1), 6188. doi: 10.1038/s41598-018-24381-4

Batailler, M., Derouet, L., Butruille, L., & Migaud, M. (2016). Sensitivity to the photoperiod and potential migratory features of neuroblasts in the adult sheep hypothalamus. Brain Struct Funct, 221(6), 3301–3314. doi: 10.1007/s00429-015-1101-0

Bolborea M, & Dale N. (2013). Hypothalamic tanycytes: potential roles in the control of feeding and energy balance. Trends Neurosci, 36(2), 91–100.

Bolborea, M., Pollatzek, E., Benford, H., Sotelo-Hitschfeld, T., & Dale, N. (2020). Hypothalamic tanycytes generate acute hyperphagia through activation of the arcuate neuronal network. Proc Natl Acad Sci U S A, 117(25), 14473–14481. doi: 10.1073/pnas.1919887117

Bottcher, M., Muller-Fielitz, H., Sundaram, S. M., Gallet, S., Neve, V., Shionoya, K., … Schwaninger, M. (2020). NF-kappaB signaling in tanycytes mediates inflammation-induced anorexia. Mol Metab, 39, 101022. doi: 10.1016/j.molmet.2020.101022

Bush, T. G., Savidge, T. C., Freeman, T. C., Cox, H. J., Campbell, E. A., Mucke, L., … Sofroniew, M. V. (1998). Fulminant jejuno-ileitis following ablation of enteric glia in adult transgenic mice. Cell, 93(2), 189–201. doi: 10.1016/s0092-8674(00)81571-8

Butruille, L., Batailler, M., Mazur, D., Prevot, V., & Migaud, M. (2018). Seasonal reorganization of hypothalamic neurogenic niche in adult sheep. Brain Struct Funct, 223(1), 91–109. doi: 10.1007/s00429-017-1478-z

Campbell, J. N., Macosko, E. Z., Fenselau, H., Pers, T. H., Lyubetskaya, A., Tenen, D., … Tsai, L. T. (2017). A molecular census of arcuate hypothalamus and median eminence cell types. Nat Neurosci, 20(3), 484–496. doi: 10.1038/nn.4495

Chaker, Z., George, C., Petrovska, M., Caron, J. B., Lacube, P., Caille, I., & Holzenberger, M. (2016). Hypothalamic neurogenesis persists in the aging brain and is controlled by energy-sensing IGF-I pathway. Neurobiol Aging, 41, 64–72. doi: 10.1016/j.neurobiolaging.2016.02.008

Chang, C., Chen, Y. T., Yeh, S. D., Xu, Q., Wang, R. S., Guillou, F., … Yeh, S. (2004). Infertility with defective spermatogenesis and hypotestosteronemia in male mice lacking the androgen receptor in Sertoli cells. Proc Natl Acad Sci U S A, 101(18), 6876–6881. doi: 10.1073/pnas.0307306101

Chauvet, N., Privat, A., & Alonso, G. (1996). Aged median eminence glial cell cultures promote survival and neurite outgrowth of cocultured neurons. Glia, 18(3), 211–223. doi: 10.1002/(SICI)1098-1136(199611)18:3<211::AID-GLIA5>3.0.CO;2-1

Chen, R., Wu, X., Jiang, L., & Zhang, Y. (2017). Single-Cell RNA-Seq Reveals Hypothalamic Cell Diversity. Cell Rep, 18(13), 3227–3241. doi: 10.1016/j.celrep.2017.03.004

Clasadonte, J., & Prevot, V. (2018). The special relationship: glia-neuron interactions in the neuroendocrine hypothalamus. [Review]. Nat Rev Endocrinol, 14(1), 25–44. doi: 10.1038/nrendo.2017.124

Culler, M. D., & Negro-Vilar, A. (1986). Evidence that pulsatile follicle-stimulating hormone secretion is independent of endogenous luteinizing hormone-releasing hormone. Endocrinology, 118(2), 609–612. doi: 10.1210/endo-118-2-609

Culler, M. D., & Negro-Vilar, A. (1987). Pulsatile follicle-stimulating hormone secretion is independent of luteinizing hormone-releasing hormone (LHRH): pulsatile replacement of LHRH bioactivity in LHRH-immunoneutralized rats. Endocrinology, 120(5), 2011–2021. doi: 10.1210/endo-120-5-2011

Deacon, R. M. (2006). Digging and marble burying in mice: simple methods for in vivo identification of biological impacts. Nat Protoc, 1(1), 122–124. doi: 10.1038/nprot.2006.20

Derouiche, L., Keller, M., Duittoz, A. H., & Pillon, D. (2015). Developmental exposure to Ethinylestradiol affects transgenerationally sexual behavior and neuroendocrine networks in male mice. Sci Rep, 5, 17457. doi: 10.1038/srep17457

Dimakopoulou, A., Foran, D., Jayasena, C. N., & Minhas, S. (2020). Stimulation of Leydig and Sertoli cellular secretory function by anti-oestrogens: Tamoxifen. Curr Pharm Des. doi: 10.2174/1381612826666200213095228

Doetsch, F., Caille, I., Lim, D. A., Garcia-Verdugo, J. M., & Alvarez-Buylla, A. (1999). Subventricular zone astrocytes are neural stem cells in the adult mammalian brain. Cell, 97(6), 703–716.

Garcia, A. D., Doan, N. B., Imura, T., Bush, T. G., & Sofroniew, M. V. (2004). GFAP-expressing progenitors are the principal source of constitutive neurogenesis in adult mouse forebrain. Nat Neurosci, 7(11), 1233–1241. doi: 10.1038/nn1340

Gibson, M. J., Ingraham, L., & Dobrjansky, A. (2000). Soluble factors guide gonadotropin-releasing hormone axonal targeting to the median eminence. [Research Support, U.S. Gov’t, P.H.S.]. Endocrinology, 141(9), 3065–3071. doi: 10.1210/endo.141.9.7656

Glover, L. R., Schoenfeld, T. J., Karlsson, R. M., Bannerman, D. M., & Cameron, H. A. (2017). Ongoing neurogenesis in the adult dentate gyrus mediates behavioral responses to ambiguous threat cues. [Comparative Study]. PLoS Biol, 15(4), e2001154. doi: 10.1371/journal.pbio.2001154

Goodman, T., Nayar, S. G., Clare, S., Mikolajczak, M., Rice, R., Mansour, S., … Hajihosseini, M. K. (2020). Fibroblast growth factor 10 is a negative regulator of postnatal neurogenesis in the mouse hypothalamus. Development, 147(13). doi: 10.1242/dev.180950

Haan, N., Goodman, T., Najdi-Samiei, A., Stratford, C. M., Rice, R., El Agha, E., … Hajihosseini, M. K. (2013). Fgf10-expressing tanycytes add new neurons to the appetite/energy-balance regulating centers of the postnatal and adult hypothalamus. The Journal of neuroscience : the official journal of the Society for Neuroscience, 33(14), 6170–6180. doi: 10.1523/JNEUROSCI.2437-12.2013

Hall, J. E., Brodie, T. D., Badger, T. M., Rivier, J., Vale, W., Conn, P. M., … Crowley, W. F., Jr. (1988). Evidence of differential control of FSH and LH secretion by gonadotropin-releasing hormone (GnRH) from the use of a GnRH antagonist. J Clin Endocrinol Metab, 67(3), 524–531. doi: 10.1210/jcem-67-3-524

Hamernik, D. L., Crowder, M. E., Nilson, J. H., & Nett, T. M. (1986). Measurement of messenger ribonucleic acid for luteinizing hormone beta-subunit, alpha-subunit, growth hormone, and prolactin after hypothalamic pituitary disconnection in ovariectomized ewes. Endocrinology, 119(6), 2704–2710. doi: 10.1210/endo-119-6-2704

Huang, L., DeVries, G. J., & Bittman, E. L. (1998). Photoperiod regulates neuronal bromodeoxyuridine labeling in the brain of a seasonally breeding mammal. J Neurobiol, 36(3), 410–420.

Imura, T., Kornblum, H. I., & Sofroniew, M. V. (2003). The predominant neural stem cell isolated from postnatal and adult forebrain but not early embryonic forebrain expresses GFAP. J Neurosci, 23(7), 2824–2832.

Kameda, Y., Arai, Y., & Nishimaki, T. (2003). Ultrastructural localization of vimentin immunoreactivity and gene expression in tanycytes and their alterations in hamsters kept under different photoperiods. Cell and tissue research, 314(2), 251–262. doi: 10.1007/s00441-003-0789-y

Knobil, E. (1990). The GnRH pulse generator. [Research Support, Non-U.S. Gov’t Research Support, U.S. Gov’t, P.H.S.]. Am J Obstet Gynecol, 163(5 Pt 2), 1721–1727. doi: 10.1016/0002-9378(90)91435-f

Kokoeva MV, Yin H, & Flier JS. (2007). Evidence for constitutive neural cell proliferation in the adult murine hypothalamus. J Comp Neurol, 505(2), 209–220.

Kokoeva, M. V., Yin, H., & Flier, J. S. (2005). Neurogenesis in the hypothalamus of adult mice: potential role in energy balance. Science, 310(5748), 679–683. doi: 10.1126/science.1115360

Lee DA, & Blackshaw S. (2012). Functional implications of hypothalamic neurogenesis in the adult mammalian brain. Int J Dev Neurosci, 30(8), 615–621.

Lee, D. A., Bedont, J. L., Pak, T., Wang, H., Song, J., Miranda-Angulo, A., … Blackshaw, S. (2012). Tanycytes of the hypothalamic median eminence form a diet-responsive neurogenic niche. Nature neuroscience, 15(5), 700–702. doi: 10.1038/nn.3079

Li, J., Tang, Y., & Cai, D. (2012). IKKbeta/NF-kappaB disrupts adult hypothalamic neural stem cells to mediate a neurodegenerative mechanism of dietary obesity and pre-diabetes. [Research Support, N.I.H., Extramural Research Support, Non-U.S. Gov’t]. Nature cell biology, 14(10), 999–1012. doi: 10.1038/ncb2562

Li, J., Tang, Y., Purkayastha, S., Yan, J., & Cai, D. (2014). Control of obesity and glucose intolerance via building neural stem cells in the hypothalamus. Mol Metab, 3(3), 313–324. doi: 10.1016/j.molmet.2014.01.012

Lincoln, G. A., & Fraser, H. M. (1979). Blockade of episodic secretion of luteinizing hormone in the ram by the administration of antibodies to luteinizing hormone releasing hormone. Biol Reprod, 21(5), 1239–1245. doi: 10.1095/biolreprod21.5.1239

Marshall, G. P., 2nd, Reynolds, B. A., & Laywell, E. D. (2007). Using the neurosphere assay to quantify neural stem cells in vivo. [Review]. Curr Pharm Biotechnol, 8(3), 141–145. doi: 10.2174/138920107780906559

Meirsman, A. C., Le Merrer, J., Pellissier, L. P., Diaz, J., Clesse, D., Kieffer, B. L., & Becker, J. A. (2016). Mice Lacking GPR88 Show Motor Deficit, Improved Spatial Learning, and Low Anxiety Reversed by Delta Opioid Antagonist. Biol Psychiatry, 79(11), 917–927. doi: 10.1016/j.biopsych.2015.05.020

Messager, S., Chatzidaki, E. E., Ma, D., Hendrick, A. G., Zahn, D., Dixon, J., … Aparicio, S. A. (2005). Kisspeptin directly stimulates gonadotropin-releasing hormone release via G protein-coupled receptor 54. Proc Natl Acad Sci U S A, 102(5), 1761–1766. doi: 10.1073/pnas.0409330102

Messina, A., Langlet, F., Chachlaki, K., Roa, J., Rasika, S., Jouy, N., … Prevot, V. (2016). A microRNA switch regulates the rise in hypothalamic GnRH production before puberty. Nat Neurosci, 19(6), 835–844. doi: 10.1038/nn.4298

Migaud, M., Batailler, M., Pillon, D., Franceschini, I., & Malpaux, B. (2011). Seasonal changes in cell proliferation in the adult sheep brain and pars tuberalis. Journal of biological rhythms, 26(6), 486–496. doi: 10.1177/0748730411420062

Migaud, M., Batailler, M., Segura, S., Duittoz, A., Franceschini, I., & Pillon, D. (2010). Emerging new sites for adult neurogenesis in the mammalian brain: a comparative study between the hypothalamus and the classical neurogenic zones. The European journal of neuroscience, 32(12), 2042–2052. doi: 10.1111/j.1460-9568.2010.07521.x

Mohr, M. A., DonCarlos, L. L., & Sisk, C. L. (2017). Inhibiting Production of New Brain Cells during Puberty or Adulthood Blunts the Hormonally Induced Surge of Luteinizing Hormone in Female Rats. eNeuro, 4(5). doi: 10.1523/ENEURO.0133-17.2017

Morshead, C. M., Garcia, A. D., Sofroniew, M. V., & van Der Kooy, D. (2003). The ablation of glial fibrillary acidic protein-positive cells from the adult central nervous system results in the loss of forebrain neural stem cells but not retinal stem cells. Eur J Neurosci, 18(1), 76–84. doi: 10.1046/j.1460-9568.2003.02727.x

Mullier A, Bouret SG, Prevot V, & Dehouck B. (2010). Differential distribution of tight junction proteins suggests a role for tanycytes in blood-hypothalamus barrier regulation in the adult mouse brain. J Comp Neurol, 518(7), 943–962.

Orgeur, P., Bernard, S., Naciri, M., Nowak, R., Schaal, B., & Levy, F. (1999). Psychobiological consequences of two different weaning methods in sheep. Reprod Nutr Dev, 39(2), 231–244. doi: 10.1051/rnd:19990208

Osterstock, G., El Yandouzi, T., Romano, N., Carmignac, D., Langlet, F., Coutry, N., … Mery, P. F. (2014). Sustained alterations of hypothalamic tanycytes during posttraumatic hypopituitarism in male mice. Endocrinology, 155(5), 1887–1898. doi: 10.1210/en.2013-1336

Parkash, J., Messina, A., Langlet, F., Cimino, I., Loyens, A., Mazur, D., … Giacobini, P. (2015). Semaphorin7A regulates neuroglial plasticity in the adult hypothalamic median eminence. Nat Commun, 6, 6385. doi: 10.1038/ncomms7385

Pavlou, S. N., Debold, C. R., Island, D. P., Wakefield, G., Rivier, J., Vale, W., & Rabin, D. (1986). Single subcutaneous doses of a luteinizing hormone-releasing hormone antagonist suppress serum gonadotropin and testosterone levels in normal men. J Clin Endocrinol Metab, 63(2), 303–308. doi: 10.1210/jcem-63-2-303

Pellegrino, G., Trubert, C., Terrien, J., Pifferi, F., Leroy, D., Loyens, A., … Sharif, A. (2018). A comparative study of the neural stem cell niche in the adult hypothalamus of human, mouse, rat and gray mouse lemur (Microcebus murinus). J Comp Neurol, 526(9), 1419–1443. doi: 10.1002/cne.24376

Pencea V, Bingaman KD, Wiegand SJ, & Luskin MB. (2001). Infusion of brain-derived neurotrophic factor into the lateral ventricle of the adult rat leads to new neurons in the parenchyma of the striatum, septum, thalamus, and hypothalamus. J Neurosci, 21(17), 6706–6717.

Pierce, A. A., & Xu, A. W. (2010). De novo neurogenesis in adult hypothalamus as a compensatory mechanism to regulate energy balance. The Journal of neuroscience : the official journal of the Society for Neuroscience, 30(2), 723–730. doi: 10.1523/JNEUROSCI.2479-09.2010

Pineda, J. L., Lee, B. C., Spiliotis, B. E., Vale, W., Rivier, J., Brown, T. J., & Bercu, B. B. (1983). Effect of GnRH antagonist, [Ac-delta 3Pro1, pFDPhe2, DTrp3,6] GnRH, on pulsatile gonadotrop in secretion in the castrate male primate. J Clin Endocrinol Metab, 56(2), 420–422. doi: 10.1210/jcem-56-2-420

Platel, J. C., Gordon, V., Heintz, T., & Bordey, A. (2009). GFAP-GFP neural progenitors are antigenically homogeneous and anchored in their enclosed mosaic niche. Glia, 57(1), 66–78. doi: 10.1002/glia.20735

Prevot V, Croix D, Bouret S, Dutoit S, Tramu G, Stefano GB, & Beauvillain JC. (1999). Definitive evidence for the existence of morphological plasticity in the external zone of the median eminence during the rat estrous cycle: implication of neuro-glio-endothelial interactions in gonadotropin-releasing hormone release. Neuroscience, 94(3), 809–819.

Prevot, V., Dehouck, B., Sharif, A., Ciofi, P., Giacobini, P., & Clasadonte, J. (2018). The Versatile Tanycyte: A Hypothalamic Integrator of Reproduction and Energy Metabolism. Endocr Rev, 39(3), 333–368. doi: 10.1210/er.2017-00235

Prieto, M., Chauvet, N., & Alonso, G. (2000). Tanycytes transplanted into the adult rat spinal cord support the regeneration of lesioned axons. Exp Neurol, 161(1), 27–37. doi: 10.1006/exnr.1999.7223

Robins, S.C., Stewart, I., McNay, D.E., Taylor, V., Giachino, C., Goetz, M., … Placzek, M. (2013). α-Tanycytes of the adult hypothalamic third ventricle include distinct populations of FGF-responsive neural progenitors. Nat Commun, 4(2049).

Rodriguez, E. M., Blazquez, J. L., Pastor, F. E., Pelaez, B., Pena, P., Peruzzo, B., & Amat, P. (2005). Hypothalamic tanycytes: a key component of brain-endocrine interaction. Int Rev Cytol, 247, 89–164. doi: 10.1016/S0074-7696(05)47003-5

Seri, B., Garcia-Verdugo, J. M., McEwen, B. S., & Alvarez-Buylla, A. (2001). Astrocytes give rise to new neurons in the adult mammalian hippocampus. J Neurosci, 21(18), 7153–7160.

Sharif A, Fitzsimons C.P., & Lucassen P. (2021). Neurogenesis in the adult hypothalamus: A distinct form of structural plasticity involved in metabolic and circadian regulation, with potential relevance for human pathophysiology (Vol. 179): Elsevier B.V.

Sousa-Ferreira L, de Almeida LP, & Cavadas C. (2014). Role of hypothalamic neurogenesis in feeding regulation. Trends Endocrinol Metab, 25(2), 80–88.

Steyn, F. J., Wan, Y., Clarkson, J., Veldhuis, J. D., Herbison, A. E., & Chen, C. (2013). Development of a methodology for and assessment of pulsatile luteinizing hormone secretion in juvenile and adult male mice. Endocrinology, 154(12), 4939–4945. doi: 10.1210/en.2013-1502

Thompson, E. L., Murphy, K. G., Patterson, M., Bewick, G. A., Stamp, G. W., Curtis, A. E., … Bloom, S. R. (2006). Chronic subcutaneous administration of kisspeptin-54 causes testicular degeneration in adult male rats. Am J Physiol Endocrinol Metab, 291(5), E1074–1082. doi: 10.1152/ajpendo.00040.2006

Wei. LC, Shi. M, Chen. LW, Cao. R, Zhang. P, & Chan. YS. (2002). Nestin-containing cells express glial fibrillary acidic protein in the proliferative regions of central nervous system of postnatal developing and adult mice. Brain Res Dev Brain Res, 139(1), 9–17.

Xu, Y., Tamamaki, N., Noda, T., Kimura, K., Itokazu, Y., Matsumoto, N., … Ide, C. (2005). Neurogenesis in the ependymal layer of the adult rat 3rd ventricle. Experimental neurology, 192(2), 251–264. doi: 10.1016/j.expneurol.2004.12.021

Yoo, S., & Blackshaw, S. (2018). Regulation and function of neurogenesis in the adult mammalian hypothalamus. Prog Neurobiol, 170, 53–66. doi: 10.1016/j.pneurobio.2018.04.001

Yoo, S., Cha, D., Kim, S., Jiang, L., Cooke, P., Adebesin, M., … Blackshaw, S. (2020). Tanycyte ablation in the arcuate nucleus and median eminence increases obesity susceptibility by increasing body fat content in male mice. Glia, 68(10), 1987–2000. doi: 10.1002/glia.23817

Zhang, G., Li, J., Purkayastha, S., Tang, Y., Zhang, H., Yin, Y., … Cai, D. (2013). Hypothalamic programming of systemic ageing involving IKK-beta, NF-kappaB and GnRH. Nature, 497(7448), 211–216. doi: 10.1038/nature12143

Zhang, Y., Kim, M. S., Jia, B., Yan, J., Zuniga-Hertz, J. P., Han, C., & Cai, D. (2017). Hypothalamic stem cells control ageing speed partly through exosomal miRNAs. Nature, 548(7665), 52–57. doi: 10.1038/nature23282

