## Supplemental Figures 1 - 5 for "Selective ablation of adult GFAP-expressing tanycytes leads to hypogonadotropic hypogonadism in males"

Supplementary Figure 1

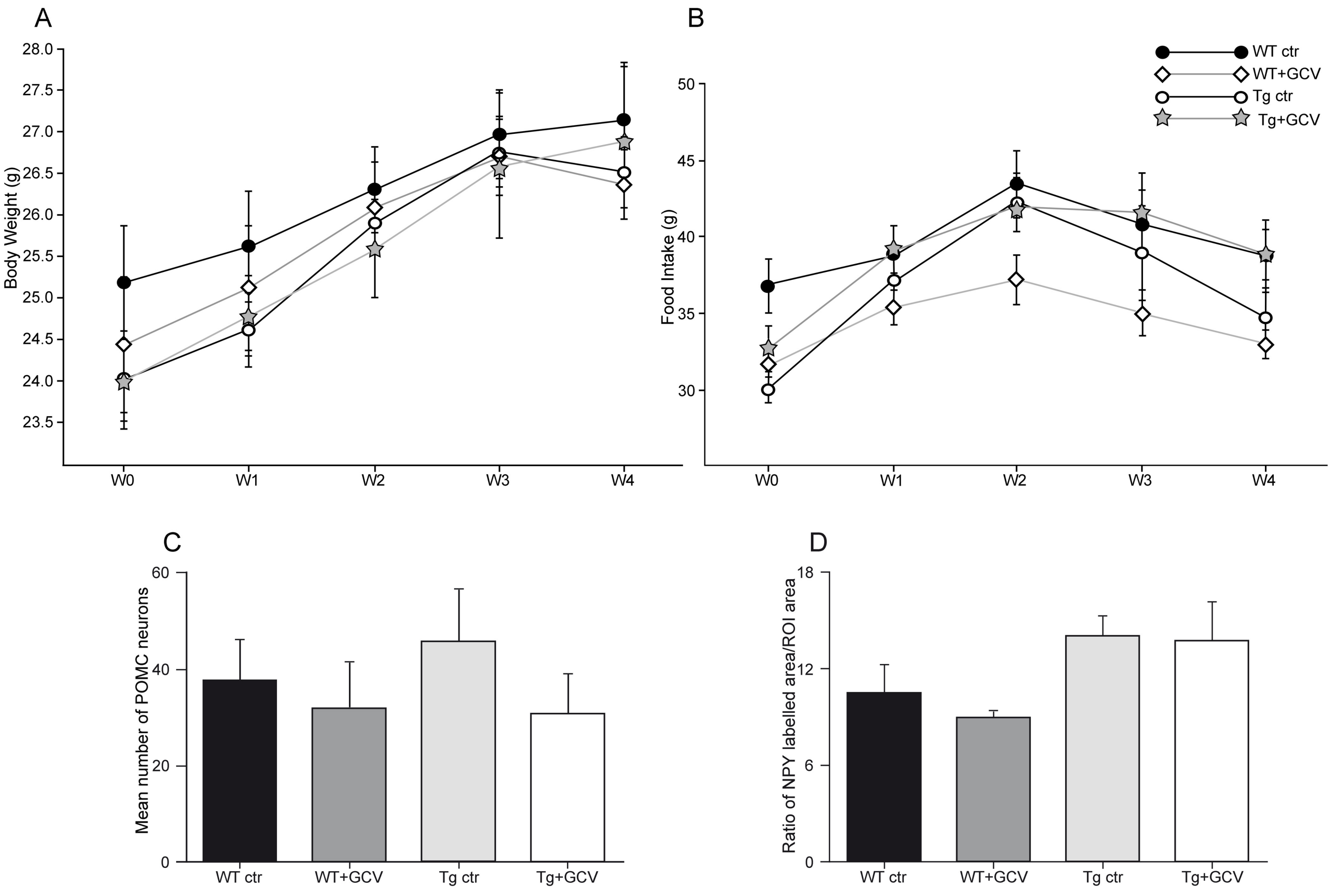


**Figure_1_Supp: Ablation of GFAP-positive tanycytes does not affect food intake**.

Body weight (A) and food intake (B) recorded during treatment are not affected in Tg+GCV male mice. The expressions of POMC (C) and NPY (D) in the arcuate nucleus are not modified by the ablation of GFAP positive NSPCs. Data are expressed as the mean±SEM, n≥5 for each group.

Supplementary Figure 2

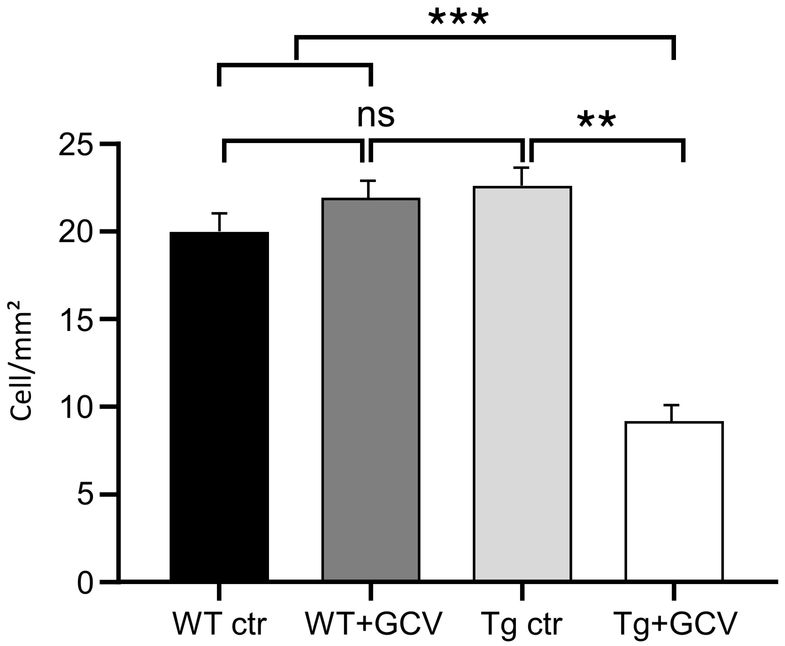


**Figure_2_Supp: Cellular analysis of the seminiferous tubules.**

The number of spermatogonia and spermatocytes per seminiferous tubule was assessed and expressed as the mean number of cells per $mm^2$ (n=3 tubules/image, 3 images/mouse and 2-3 mice/group). Data are expressed as the mean±SEM. **$p<0.01$, ***$p<0.001$.

Supplementary Figure 3

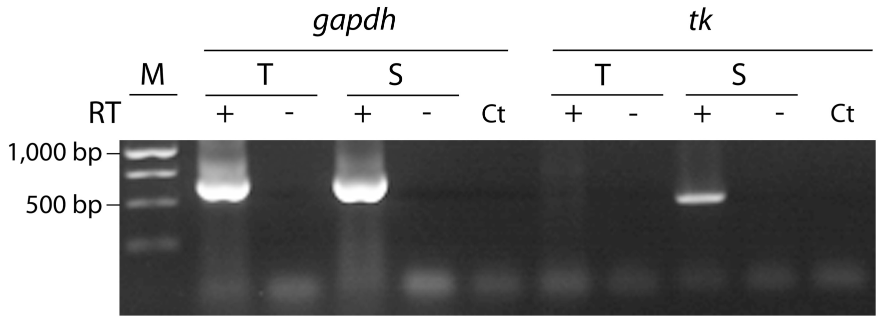


**Figure_3_Supp. Assessment of transgenic Tk mRNA expression by RT-PCR.**

The total RNA from the testis and the striatum of three Tg mice was analysed by RT-PCR. RT-PCR was performed as described in the Methods section either in the presence (+) or absence (−) of reverse transcriptase to control for DNA contamination. A PCR product of 495 bp for Tk was detected in the striatum but not in the testis. A PCR product of 619 bp for Gapdh was detected in both tissues. Gapdh, Glyceraldehyde -3 -phosphate dehydrogenase; Tk, Thymidine kinase
M, 1-kb marker; T, testis; S, striatum.

Supplementary Figure 4

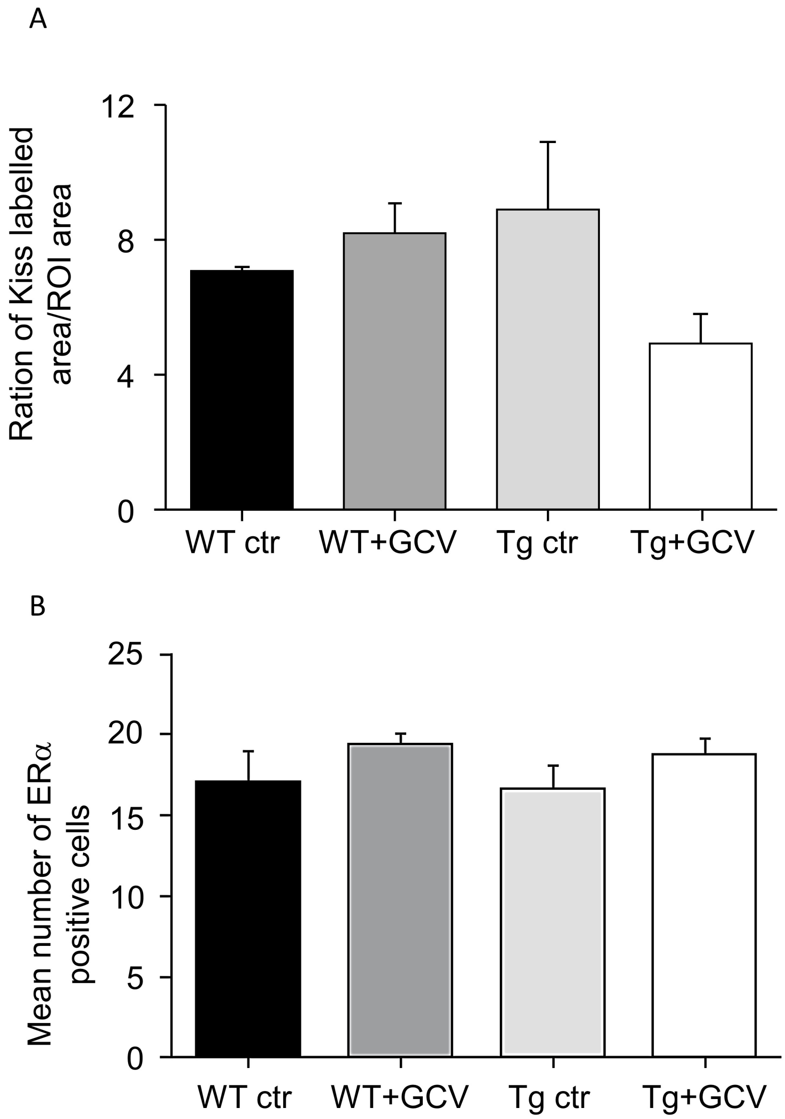


**Figure_4_Supp: Ablation of GFAP-positive tanycytes does not affect the expression of Kisspeptin and ER alpha.**

Kisspeptin labelling (A) and mean number of ERa-positive cells (B) are not affected by the ablation of GFAP-positive tanycytes. Data are expressed as the mean±SEM, n≥3 for each group.

Supplementary Figure 5

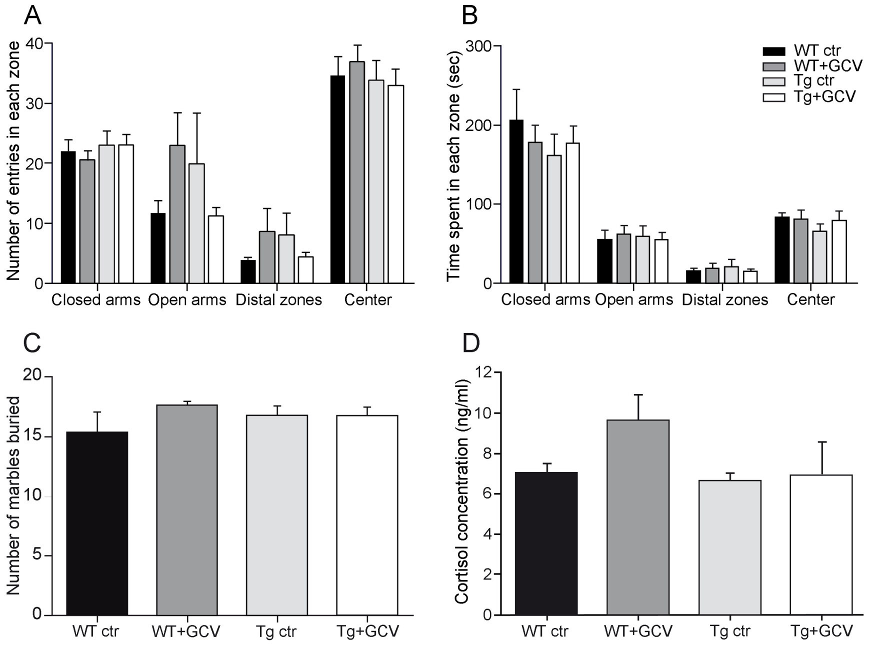


**Figure_5_Supp. Ablation of GFAP-positive tanycytes does not affect the level of anxiety.**

The numbers of entries (A) and the time spent (B) in each zone of the elevated plus maze are not affected by the ablation of GFAP-positive tanycytes. (C) The number of marbles buried in a marble burying test is identical between the four experimental groups.
(D) Tg+GCV male mice display comparable plasma cortisol concentrations than the three control groups. Data are expressed as the mean±SEM, n≥3 for each group.
